# Modelling inter-shot variability for robust temporal sub-sampling of dynamic, GABA-edited MR spectroscopy data

**DOI:** 10.1101/2024.12.05.627018

**Authors:** Alexander R. Craven, Lars Ersland, Kenneth Hugdahl, Renate Grüner

**Author notes:** **Corresponding author:** Alexander R. Craven (M.Sc.) Department of Biological and Medical Psychology, University of Bergen Jonas Lies vei 91, 5009 Bergen, Norway.

## Abstract

Variability between individual transients in an MRS acquisition presents a challenge for reliable quantification, particularly in functional scenarios where discrete subsets of the available transients may be compared. The current study aims to develop and validate a model for removing unwanted variance from GABA-edited MRS data, whilst preserving variance of potential interest – such as metabolic response to a functional task.

A linear model is used to describe sources of variance in the system: intrinsic, periodic variance associated with phase cycling and spectral editing, and abrupt changes associated with subject movement. We broadly hypothesize that modelling these factors appropriately will improve spectral quality and reduce variance in quantification outcomes, without introducing bias to the estimates. We additionally anticipate that the models will improve (or at least maintain) sensitivity to functional changes, outperforming established methods in this regard.

In vivo GABA-edited MRS data (203 subjects from the publicly available Big GABA collection) were sub-sampled strategically to assess individual components of the model, benchmarked against the uncorrected case and against established approaches such as spectral improvement by Fourier thresholding (SIFT). Changes in metabolite concentration and lineshape simulating response to a functional task were synthesized, and sensitivity to such changes was assessed.

Composite models yielded improved SNR and reduced variability of GABA+ estimates compared to the uncorrected case in all scenarios, with performance for individual model components varying. Similarly, while some model components in isolation led to increased variability in estimates, no bias was observed in these or in the composite models. While SIFT yielded the greatest reductions in unwanted variance, the resultant data were substantially less sensitive to synthetic functional changes.

We conclude that the modelling presented is effective at reducing unwanted variance, whilst retaining temporal dynamics of interest for functional MRS applications, and recommend its inclusion in fMRS processing pipelines.

**Highlights:**

- A novel technique for modelling unwanted variance between transients is investigated.
- Suitable covariate models yield improved SNR and reduced variability in GABA+ estimates from the resultant spectra.
- Extracted spectra remain sensitive to temporal dynamics of interest for functional MRS applications.

**Graphical Abstract:** In dynamic MRS analysis, unwanted variability between transients may confound findings when sub-sampling within a single acquisition. We investigate covariate models and lineshape matching strategies to address this. We present composite models yielding improved quality metrics and within-scan repeatability while maintaining sensitivity to (synthetic) functional changes.

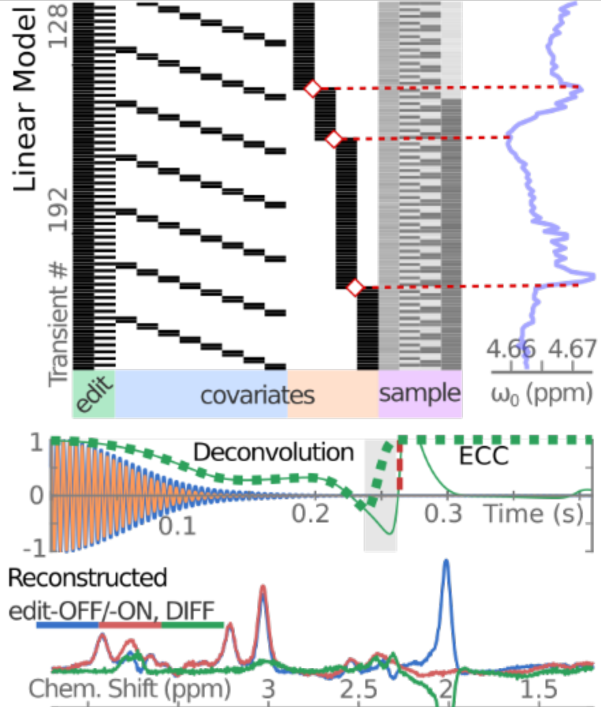

## 1 Introduction

Magnetic Resonance Spectroscopy (MRS) is an inherently noisy technique, with variability between individual transients presenting an ongoing challenge for reliable quantification. Indeed, this is the primary motivation for acquiring multiple transients – with the expectation that unwanted variability will either cancel out (destructive interference) or be averaged away to insignificance. However, in applications such as functional MRS (fMRS) where transients may be binned, combined and compared across different parts of an acquisition, it is not clear that the underlying assumptions will hold. This may in turn lead to less reliable estimates and potentially biased outcomes. We therefore propose explicitly modelling inter-transient variability, and subtracting the resultant components to yield more robust spectral estimates when quantifying a subset of the available transients.

Variability between transients in an MRS experiment may be broadly discussed in terms of its origins and its temporal characteristics: whether it is intrinsic to the sequence, characteristic of the hardware, or reflective of in vivo physiological processes, and whether it presents as periodic variation, slow variation, a sudden change, or a stochastic process (adopting the terminology of Hennig 1991^1^).

Most PRESS localized MRS acquisitions incorporate a phase cycling scheme^1–6^ for coherence pathway selection, either relying on cancellation of unwanted signals when averaged across the full cycle, or adopting a Fourier-transform approach to isolate the proper signal^1^. Spectral editing techniques such as MEGA-PRESS^7,8^, and other advanced techniques like metabolite cycling, introduce additional cyclic elements which are integral to the acquisition, and generally perform best when data are balanced across a full cycle; imperfect cancellation will often give rise to unwanted signals and artefacts in the resultant spectrum. If transients are sampled in a scheme which does not align to all these cyclic elements, such artefacts are likely to arise. As well as these intrinsic cyclic factors, *periodic signal variations* may also arise from physiological processes – respiratory and circulatory factors, for example – which are generally not aligned to the sequence-intrinsic elements.

In the context of dynamic or functional MRS, the design of the task (for example, distribution of stimuli into task-ON, task-OFF blocks) may introduce an additional *periodic* factor. Furthermore, changes associated with a functional task may occur on different timescales which may lag the stimulus: blood-oxygenation-related changes in local T2/T2* properties on a timescale described by the blood oxygenation level dependent (BOLD) haemodynamic response function (HRF) (typically 0-10 seconds), metabolic changes on a shorter timescale (perhaps in the order of hundreds of ms)^9,10^, and incidental subject movement (for example, hand movement or vocalisation in response to the task).

Temperature-related field drift^11^ and gradual changes in physiological state may give rise to *slow signal variations*, which can lead to accumulating cancellation errors as well as broadening of the resultant spectra. For spectral editing techniques, frequency drift may also degrade editing efficiency, as the selective refocussing pulses drift away from the nominal target frequency. In a sub-sampling or windowing scheme where conditions are compared sequentially (for example, task-OFF blocks followed by task-ON blocks in a functional design), one side of the contrast may systematically experience greater effects of accumulated drift.

*Sudden signal changes* are often the result of patient motion during the examination. Although generally occurring at unpredictable intervals, they are often identifiable by abrupt changes in the apparent centre frequency of residual water. Real-time frequency adjustment and other periodic navigators present a special case, as they may introduce sudden changes at regular intervals, to compensate for gradual changes.

Finally, there are *stochastic signal variations*, such as RF noise and Johnson-Nyquist noise in the receiver system. Stimuli in an event-related fMRS paradigm may also present as stochastic variations on a longer time scale.

Various strategies exist for dealing with variability in MRS data; advanced spectral registration techniques can mitigate some issues relating to field drifts and shifts, while techniques such as Rank-Order Statistical Filtering (OSF)^12^ and Spectral Improvement by Fourier Thresholding (SIFT)^13^ have been demonstrated to improve signal-to-noise ratio (SNR) and reduce variability in the general case. However, these techniques may not be optimal for fMRS applications: OSF typically collapses (or un-orders) the temporal dimension, while SIFT explicitly filters weaker periodic components in the frequency domain. Although the SIFT technique has been applied successfully in functional contexts for ^31^P spectra^14^, it relies on judiciously-chosen thresholds to distinguish between components of interest and nuisance signals. While this has been shown to be effective for random variations, there is no guarantee that subtle changes in metabolite levels will outweigh periodic signal variations associated with phase cycling for example, and it is also not clear whether the technique will be suitable for randomly distributed variation of interest in an event-related paradigm.

The present study investigates a linear modelling technique for separating variability of interest (for example, related to a functional task) from variability associated with nuisance factors, as predicted by intrinsic factors in the sequence (e.g., phase cycling) and inferred subject motion. We evaluate this model in terms of signal quality and within-acquisition test-retest reliability on in vivo data acquired at rest (baseline), and sensitivity to change in relation to a simulated functional task, comparing against both the uncorrected data and spectra filtered with the more established SIFT technique.

## 2 Methods

### 2.1 In vivo source data

The present study focusses on data acquired in vivo with the MEGA-PRESS^7,8^ editing sequence, for detection of γ-aminobutyric acid (GABA). The GABA peak is modelled together with underlying co-edited signals, and reported as GABA+.

GABA+ data from 203 consenting adult volunteers (age 18-36 years, approximately even female/male split, reporting no known neurological or psychiatric illness) were obtained from the publicly available Big GABA collection of datasets^15,16^, originally acquired in an international collaborative study. These data originated from sixteen 3T MRI scanners from three major manufacturers (GE, Philips, Siemens), each at a different site. GABA-edited spectra (GABA+: TR/TE=2000/68 ms, 320 transients, with editing pulses at 1.9 and 7.46 ppm for edit-ON/-OFF respectively) had been obtained from a 3×3×3 cm^3^ voxel in the posterior cingulate region while the subjects were at rest (i.e., without a functional task). These data were collected in accordance with ethical standards of the respective local institutional review boards (IRB), with subjects explicitly consenting to the sharing of anonymized data for further studies.

This collection of datasets has been extensively characterised and reported in a number of published studies ^15–17^; full details on sample demographics, acquisition protocol, software and hardware configurations may be found in these papers; vendor-specific factors have also been examined previously. A summary of pertinent details is reproduced in Supplementary Table 1 (MRSinMRS^18^ checklist) and Supplementary Table 2.

Original data in vendor-specific format were imported using the GannetLoad function from Gannet (v3.1); an initial fit was performed using the default pipeline and parameters, to yield preliminary estimates on GABA+ concentration, linewidth, water signal area and water frequency which were used to inform subsequent steps. Individual transients, after eddy current compensation^19^ and robust spectral registration^20^ but without Gannet-default line-broadening or zero-fill, were taken for further modelling (see sections 2.2 and 2.3). After this modelling, the standard Gannet GABA+Glx fit was applied, with GABA+ estimates scaled according to the modelled water area, without accounting for tissue composition within the voxel: all reported findings are based on proportional change relative to an initial estimate, making further adjustments unnecessary.

### 2.2 Modelling temporal variability

Covariates for phase cycling and inferred subject motion were incorporated into a linear model for spectral combination, as illustrated in Figure 1a and described in a previous work^21^. Briefly, the model of the form *Y=XB+U* expresses each transient Y_n,*_ (pointwise, on the frequency-domain spectrum) as a combination of a nominal edit-OFF sub-spectrum (factor X_*,1_) and a periodic component associated with editing (factor X_*,2_ characterising the difference spectrum, DIFF), with parameter matrix B containing spectra modelled to the observations and U containing modelling residuals (uncorrelated noise). Additional nuisance components for periodic phase cycling (factors X_*,2+(1:n_phase)_) and partitioning according to inferred subject motion (factors X_*,2+n_phase+(1:n_motion)_) are incorporated, as detailed below.

**Figure 1.**
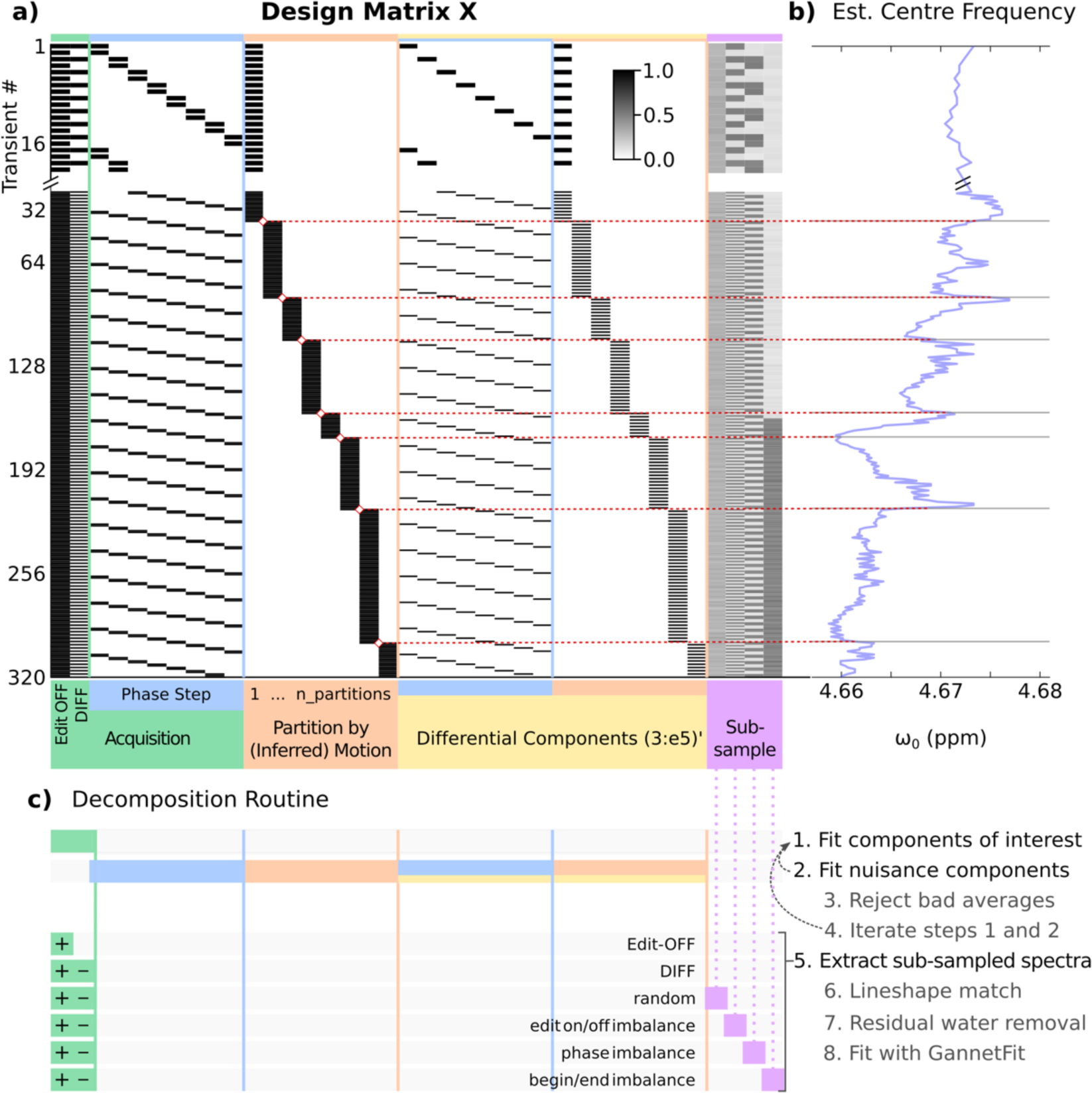
Covariate model and sampling schemes (a), with the motion covariate inferred from abrupt changes in the centre frequency estimate (b). The system is solved piecewise (c). Sub-sampling masks presented rightmost in (a) are probabilistic, derived from the average binary mask across all repetitions for all subjects. Figure derived from an earlier work^21^.

The model is solved piecewise, iteratively: after an initial fit with only the components of interest (edit-OFF and DIFF), nuisance factors are modelled to the residuals and removed, before revising the estimate for the main fit and repeating until the relative improvement drops below a defined threshold. This modelling is performed on the full set of transients, before sub-sampling of transients with the modelled nuisance components removed.

#### 2.2.1 Intrinsic cyclic effects

Phase cycling and rotation techniques^1,22^ aim to isolate signals from the desired coherence pathway, while suppressing signal from unwanted coherences (perhaps originating outside the prescribed volume of interest, or a result of imperfect flip angles). Phase of the RF pulses and receiver are shifted systematically, cyclically throughout the acquisition, such that unwanted coherences interfere destructively and cancel. Incomplete or unbalanced sampling across the cycle will impede effective cancellation of these coherences; this may be exacerbated by subject motion. Figure 2a illustrates variability across the phase cycle for a single subject – note divergence to the left of the major peaks (around 3.2 and 3.9 ppm), and in the background signal between these peaks (3.4 to 3.6 ppm), sufficient to perturb the baseline model. These effects are generally mild in the present dataset, although severity will depend on voxel placement and slice order. In the current model, unwanted coherences at each step in the phase cycle are modelled with a series of factors X_*,2+(1:n_phase)_.

**Figure 2.**
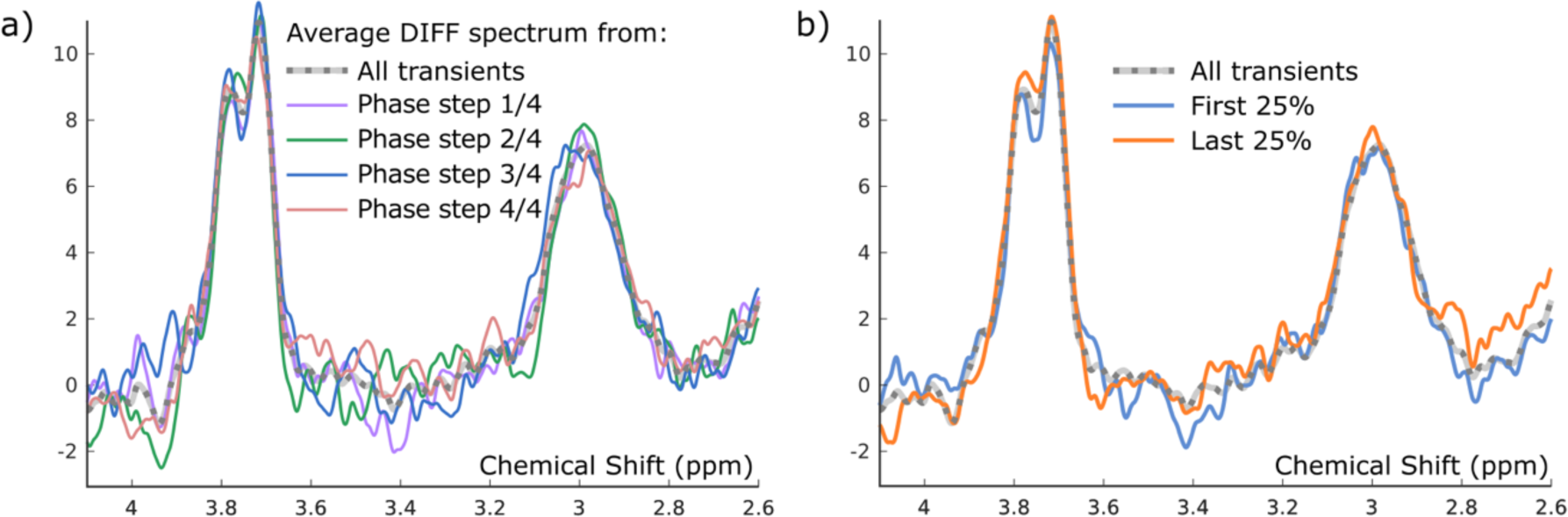
Illustrative variability in spectral data (single subject, DIFF spectrum, after spectral registration) taken from different steps in the phase cycle (a) and from earlier and later parts of the acquisition (b), the latter illustrating degraded shim from cumulative effects of subject motion and field drift throughout the scan. Differences in peak shape and background signal are apparent in each.

#### 2.2.2 Inferred subject motion

In addition to subtly changing the coverage of the spectroscopy voxel, movement of the subject is likely to induce changes in the overall shape of the acquired spectrum, including altered background, lineshape and frequency shift through variation of the local shim conditions. This variability is illustrated in Figure 2b. In the present study, we use sudden changes in the observed water frequency (illustrated in Figure 1b) to infer times at which the subject may have moved, or at which some other occurrence gave rise to a change in the acquisition conditions. Such changes are detected using the `findchangepts` function in Matlab ^23^ (R2021a), as detailed previously^21^. The set of transients is partitioned about each of these identified change points, defining model factors X_*,2+n_phase+(1:n_motion)_). Although we do not explicitly model thermal drift, the temporal partitioning performed here additionally affords the model flexibility to compensate for some drift-related changes over time.

### 2.3 Mitigating lineshape variability

Changing scan conditions, whether due to subject motion, field drift, or BOLD-related changes in response to a functional task, may alter the spectral lineshape. This may in turn impact the spectral modelling and quantification^24^. Further to the linear modelling approach for handling covariates, several strategies for modelling and compensating for lineshape changes are also assessed.

#### 2.3.1 Lineshape matching by deconvolution

In addition to simply incorporating inferred motion components as regressors of no interest in our linear model, we also consider the efficacy of lineshape matching across different sections of the acquired data – using the same partitioning scheme as for motion compensation. This is achieved using an adaption of well-established deconvolution techniques ^25–27^, combining reference deconvolution ^28^ with a crossover to ECC ^29^. The basic deconvolution is performed as a division of time-domain free induction decay (FID) signals (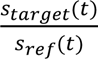, see Figure 3c), with the target lineshape *s_target_(t)* determined from the average edit-OFF spectra across the entire scan (Figure 3a), filtered to include only the primary Creatine peak around 3.0 ppm (Figure 3b), after subtracting a linear baseline along that range. The reference signal *s_ref_(t)* is obtained by applying the same procedure to each set of transients in the partitioned spectrum. A crossover point to ECC was determined on a per-reference basis, by identifying discontinuities (vertical asymptotes) in the quotient indicative of excessively small values in the denominator as the FID signal decays (Figure 3c). The resulting transformation is applied individually to each transient in the corresponding partition (both edit-ON and edit-OFF); see Figure 3d.

**Figure 3.**
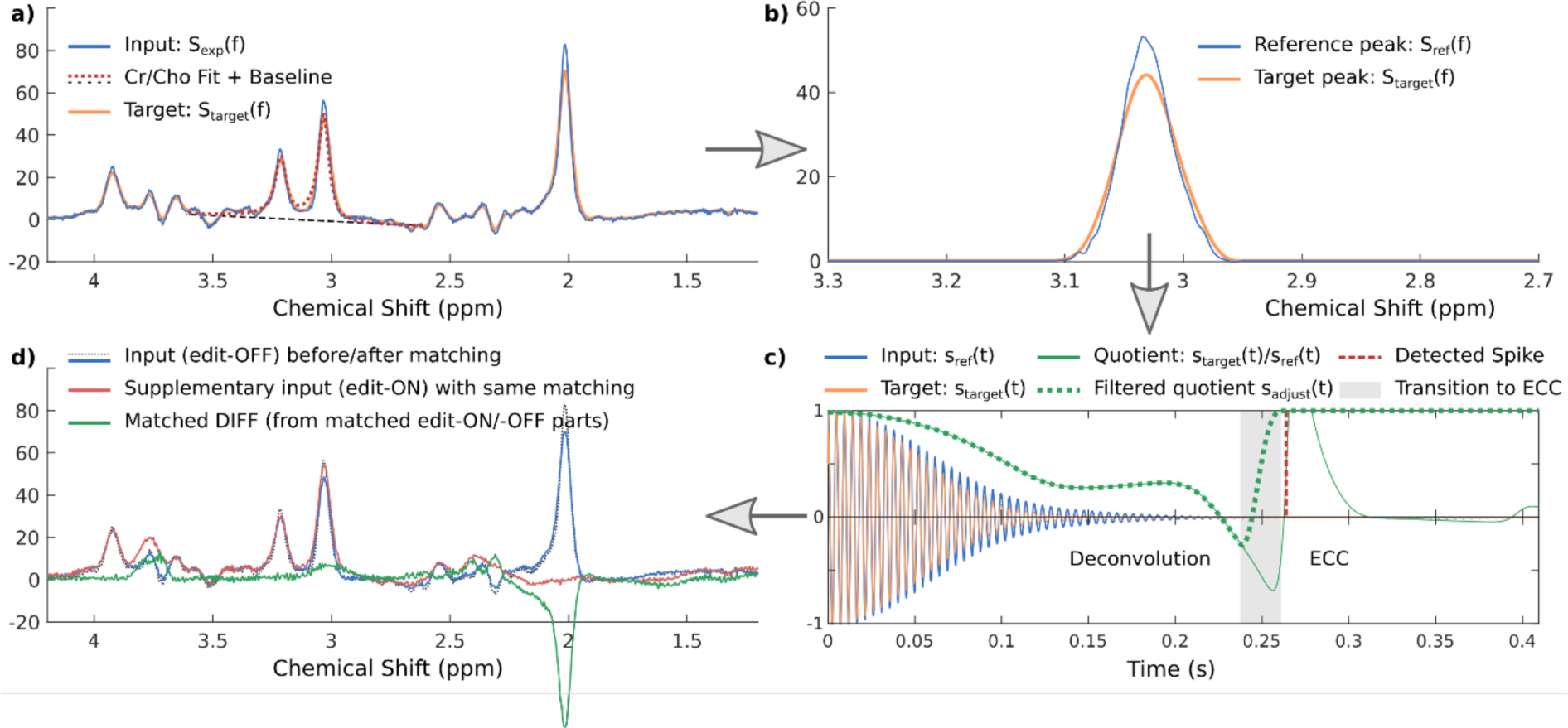
Lineshape matching by reference deconvolution: a Cho/Cr peak model on the incoming edit-OFF spectra (a) is used to delineate the Cr reference peak (b) as the basis for the deconvolution (c). The resulting transformation is applied to all transients, edit-OFF and edit-ON (d)

#### 2.3.2 Linewidth matching with a broadening function

The following lineshape matching variants are performed on individual transients, *before* the linear modelling described in section 2.2; functional variants are only relevant to simulated functional data described in section 2.5:

- **lineshape_match_part** : for each partition (defined by inferred motion), lineshape is matched to that of the full spectrum, using reference deconvolution as described in section 2.3.1.
- **lineshape_match_func** : lineshape for functional task-ON and task-OFF transients (separately) is matched to that of the full spectrum, using reference deconvolution.
- **lineshape_match_func_hrf** : after convolving the functional block (or event) timing with a canonical HRF (normalised to magnitude 1), transients having the theoretical BOLD magnitude > 50% and those < 50% are separately matched to the full spectrum, using reference deconvolution.

The following lineshape matching variants are performed on the extracted spectra *after* the linear modelling as described in section 2.2:

- **lineshape_match_late** : lineshape matching of each resultant sub-spectrum to the complete spectrum, using reference deconvolution
- **linewidth_match_lor** : linewidth matching of each resultant sub-spectrum to the complete spectrum, with Lorentzian broadening (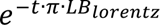).
- **linewidth_match_gau** : linewidth matching of each resultant sub-spectrum to the complete spectrum, with Gaussian broadening (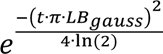)^30,31^.
- **linewidth_match_LGmix** : linewidth matching with an equal mix of Lorentzian and Gaussian broadening, with LB_lorentz_ and LB_gauss_ scaled to yield nominal linewidth in the resultant Voigt kernel.

### 2.4 Evaluating model efficacy with baseline in vivo data

For each complete spectrum, pseudo-random subsamples of the available transients are defined, targeting variation in certain scenarios given the following characteristics:

- **Random**: the sample consists of a random selection of transients, selected with equal probability across the entire sequence. This scenario characterises variance across the complete sequence.
- **Imbalanced editing**: the selected sample is comprised of 20% edit-OFF, 80% edit-ON transients. This captures unwanted variance specifically relating to the cyclic editing; we note that targeting proportionally more edit-ON than edit-OFF transients has been proposed^32^ as a strategy to bolster SNR in functional applications.
- **Imbalanced phase**: the selected sample is comprised of 20% transients from “odd” steps in the phase cycle, 80% from “even” steps, thereby emphasizing part of the variance associated with phase cycling. This may reflect a functional block design where stimulus blocks do not align to complete phase cycles.
- **Imbalanced time**: the selected sample consists of 20% of transients taken from the first half of the acquisition, and 80% taken from the last half – emphasizing variance associated with field drift and accumulated motion. In fMRS applications, this may reflect paradigms with a small number of long stimulus blocks.

Sampling patterns for each of these scenarios are illustrated in the rightmost columns of Figure 1a, where the average binary mask across subjects and repetitions shows the probability that a given transient contributes to the sample.

Each of the discrete model components (phase covariate, motion covariate, lineshape/linewidth) was assessed separately, and as composite models incorporating a combination of factors (phase + motion, or phase + motion + lineshape, for different lineshape strategies described in section 2.3). For each scan, the discrete and composite models were applied to random subsets of n=100 transients selected according to the above criteria, for 10 repetitions. The uncorrected, no-model case (simple averaging across the selected transients) is also assessed as a baseline for performance.

We additionally evaluate performance using a local implementation of the SIFT algorithm, with acceptance thresholds set to 2 and 1 standard deviations (noting that the former aligns with suggested values in the literature^13,14^).

### 2.5 Evaluating sensitivity to simulated functional changes

Time courses were defined to represent typical functional designs: an alternating block design (40 seconds rest, 20 seconds active, repeated throughout the scan to give a total of 10 task-ON blocks and 11 task-OFF blocks), and a stochastic event-related design with one third of transients randomly chosen as corresponding to active (task-ON) events, the remainder considered to be rest (task-OFF).

For transients considered as “task-ON”, a scaled, simulated metabolite component was added to the existing in vivo metabolite spectrum. The GABA component was extracted from published MEGA-PRESS basis sets distributed with the Osprey package ^33^, with vendor-specific refocusing pulses and timings – originally simulated using fast spatially resolved 2D density-matrix simulations^34^ implemented in FID-A^35^, with coefficients from Kaiser et al.^36^.

The GABA basis component was subject to line broadening to approximately match the initial estimate on in vivo GABA linewidth, and amplitude scaling relative to the initial GABA+ estimate, via the initial modelled water area estimate – all evaluated per scan.

Additive concentrations from 0 - 20 % (in steps of 3.33%) were applied to task-ON transients. Line broadening was also applied in varying degrees, to simulate BOLD-related T2/T2* effects which may be anticipated in functional spectroscopy studies^37^. Masks defining task-ON transients were convolved with a canonical HRF, then scaled to give a nominal reduction in linewidth of 0, 0.3 and 1.0 Hz in relation to task-ON periods. A 50% mixture of Lorentzian and Gaussian broadening functions was applied. To avoid possible numeric instability associated with negative linebroadening factors, this was applied relative to a fixed linebroadening of 1.2 Hz (hence, the actual applied broadening was 1.2 Hz for rest, and [1.2, 0.9, 0.2] Hz for active periods after HRF convolution).

Each discrete model component (phase, motion, lineshape per section 2.3) and composite model (phase + motion, phase + motion + lineshape) was evaluated for each level of simulated metabolite change, each level of simulated line broadening, and each basic functional design (block, event).

### 2.6 Quality control

Several rejection criteria are applied to each of the individual fitting outcomes (after any sub-sampling and modelling), with full-width at half maximum (FWHM) linewidth and Signal-to-Noise Ratio (SNR) as reported by Gannet: using GABA+ from the DIFF spectrum and NAA from the edit-OFF sub-spectrum, with SNR measured relative to detrended, downfield noise in the corresponding signal. Individual fits having excessive linewidth (FWHM_GABA+_ > 30 Hz, FWHM_NAA_ > 12 Hz), extremely low SNR (SNR_GABA+_ <5, SNR_NAA_ < 50), reported fit error GABA_FitError_W > 20%, or negative/extreme outlier values on any of these metrics (indicating failure of the fitting model) were rejected. Finally, fits with extreme outlier GABA+ estimates, deviating from the median by more than five times the Median Absolute Deviation (MAD), were rejected.

### 2.7 Numeric and statistical analysis

Analysis of quantification outcomes was performed using local scripts implemented in Python (v3.9.17), with numeric methods from the pandas ^38^ (v1.5.2), NumPy ^39^ (v1.23.5) and SciPy ^40^ (v1.9.3) packages and visualisation tools from matplotlib ^41^ (v3.3.4) and seaborn ^42^ (v0.11.2). After assessing normality (Shapiro-Wilk method ^43^) and comparability of variance (Fligner-Killeen’s test ^44^), hypothesis testing was performed with the Wilcoxon Signed Rank test ^45^. Holm-Bonferroni ^46,47^ correction is used, with adjusted p-values denoted p_holm_, and a corrected significance threshold defined as p_holm_<0.05. Where confidence intervals derived from parametric bootstrapping (10,000 permutations) are reported, these are denoted CI_95%,boot_.

Model performance was assessed using the rejection rate (section 2.6), SNR_GABA+_ and Coefficient of Variation (CV) metrics (the latter across repeated sub-samples of transients, within scan), all compared against the uncorrected, no-model (simple averaging) case. Potential bias was also assessed for each model, using the Mean Signed Difference (MSD) relative to outcomes from the full set of transients per scan, on a trimmed set of outcomes (omitting 10% of estimates from each end). Sensitivity to (synthetic) functional GABA+ changes is assessed by robust linear regression with the Theil-Sen estimator^48–50^.

## 3 Results

### 3.1 Baseline in vivo data

#### 3.1.1 Spectral quality

Reconstructed spectra for a representative subject are presented in Figure 4 for a selection of models, and Supplementary Figure 1 for the full set of models and scenarios, demonstrating reduction of spurious signal by discrete and composite model factors. Achieved SNR, linewidth, fit error and rejection rates according to the criteria in section 2.6 are presented in Supplementary Table 3, for each covariate model and each of the sampling scenarios. SNR_GABA_ differences are also presented graphically in Figure 5a, with statistical differences explored in Supplementary Table 4.

**Figure 4.**
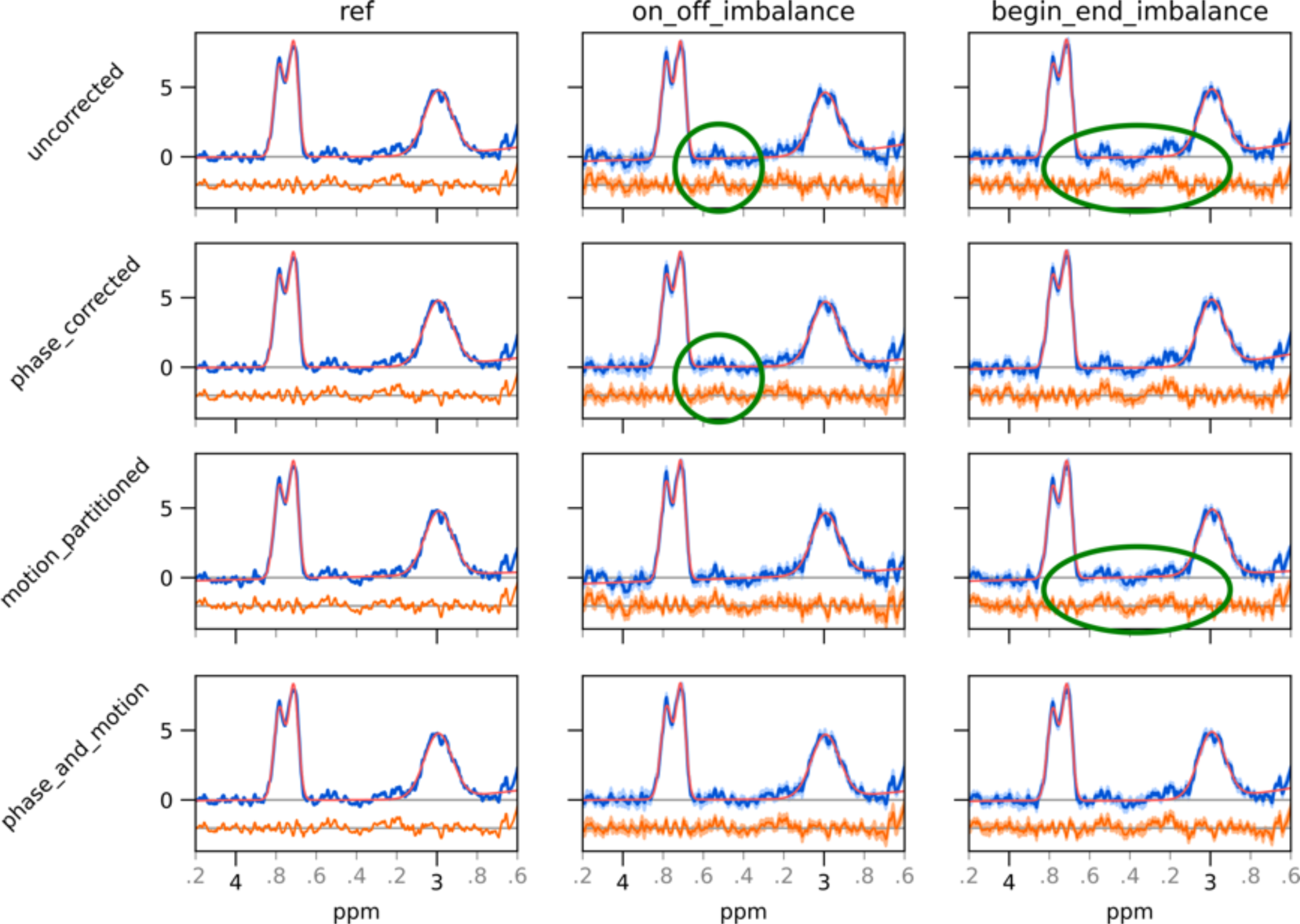
Reconstructed spectra and residuals, for a selection of sampling scenarios and modelled factors, demonstrating reduction of spurious signals (green ellipses). Mean data (blue) and residuals (orange, lower part) for a single subject are presented, with standard deviation across repetitions shaded and mean GABAGlx model fit shown in red. Additional modelling factors and sampling scenarios are presented in Supplementary Figure 1.

**Figure 5.**
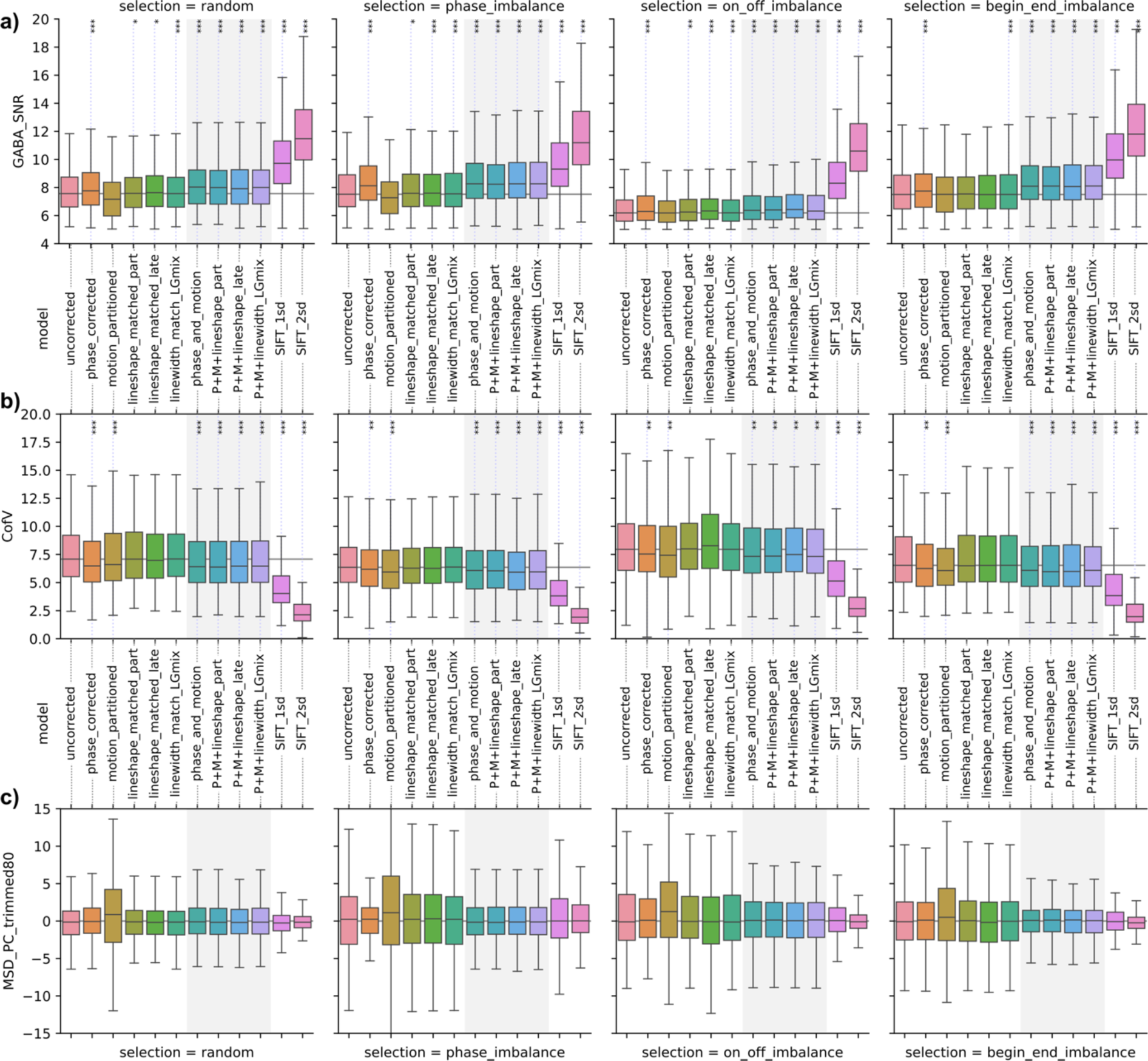
SNR (a), CV (b) and MSD (c) of GABA+ estimates from each of the models, for each scenario; significance indicated by *** p_holm_<0.001, ** p_holm_<0.01, * p_holm_<0.05

All subsampling scenarios showed reduced quality in terms of both SNR and overall rejection rates (typically ∼45% reduction in SNR_GABA_), compatible with expectations given the substantially smaller number of transients contributing to the spectra for quantification (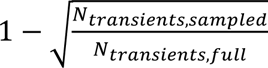= 0.44). Imbalanced sampling of edit-ON vs edit-OFF transients resulted in more substantial degradation: roughly 19% further reduction in SNR_GABA_ compared with other scenarios; this led to a high rejection rate (>20%) for spectra modelled in this sampling scenario, which did not improve with our covariate models but was ameliorated by the SIFT technique. Exploratory analysis confirmed that this pattern presents regardless of whether it is the edit-ON or the edit-OFF transients that are over-represented in the sample. For the accepted spectra, the SIFT method achieved the greatest improvements in SNR, up to 73.5% (p_holm_<0.001, CI_95%,boot_ [71.4,75.4]) for the 2 SD threshold in the imbalanced editing scenario. All composite models also yielded strongly significant improvements, albeit more modest in scale: up to 10.1% (p_holm_<0.001, CI_95%,boot_ [9.08,11.1]). Amongst composite models, those with lineshape matching of resultant sub-spectra by reference deconvolution (lineshape_match_late) generally performed marginally better than other techniques. Inclusion of the phase correction component alone gave significant improvements in all scenarios, as did most lineshape matching strategies in most scenarios. Including only the motion component in isolation did not yield any improvement. This pattern is reflected in the Fit Error, which is reduced by 38.1% (p_holm_<0.001, CI_95%,boot_ [26.6, 29.4,]) for SIFT with a 2 SD threshold, and up to 6.76% (p_holm_<0.001, CI_95%,boot_ [5.90, 7.68]) for composite models.

Additional statistics for Linewidth are presented in Supplementary Table 5 and Supplementary Figure 2. Additional statistics for GABA+ model Fit Error are presented in Supplementary Table 6 and Supplementary Figure 3.

#### 3.1.2 Test-retest reliability

Test-retest reliability within scan is summarised in Figure 5b, with full statistics in Supplementary Table 7. This shows a modest but statistically significant improvement in test-retest reliability for models incorporating the phase or motion covariates in isolation (CV reduced by up to 6.37% (p_holm_<0.001, CI_95%,boot_ [3.80, 8.82]), and lineshape components in isolation not yielding significant improvements. Composite models performed slightly better, consistently across scenarios and across lineshape strategies – with CV reduced by up to 8.06% (p_holm_<0.001, CI_95%,boot_ [5.32, 10.53]). The SIFT method resulted in more substantial improvements in test-retest reliability (25-65% reduction), for all sampling scenarios.

Mean signed deviation (MSD) was used to assess any bias introduced by the modelling; outcomes are presented in Figure 5c and Supplementary Table 8. None of the models were found to introduce statistically significant bias in the obtained concentration estimates (all p_holm_>0.05), although modelling with the motion covariate alone led to a greater spread in estimates.

### 3.2 Sensitivity to simulated functional changes

Measured response to simulated functional changes in GABA concentration, applied to vivo data, is presented in Figure 6. Additional combinations of stimulus type (block or event), simulated BOLD-related linewidth change and covariate model are presented in Supplementary Figure 4 and Supplementary Table 9. All composite models (and most discrete factors) gave similar performance to the uncorrected case in terms of coefficient of determination (R^2^), and marginally improved sensitivity to event-related stimulus, as assessed by the linear regression coefficient: typically around [0.91, 0.94, 0.97] for uncorrected, discrete model factors and composite models respectively.

**Figure 6.**
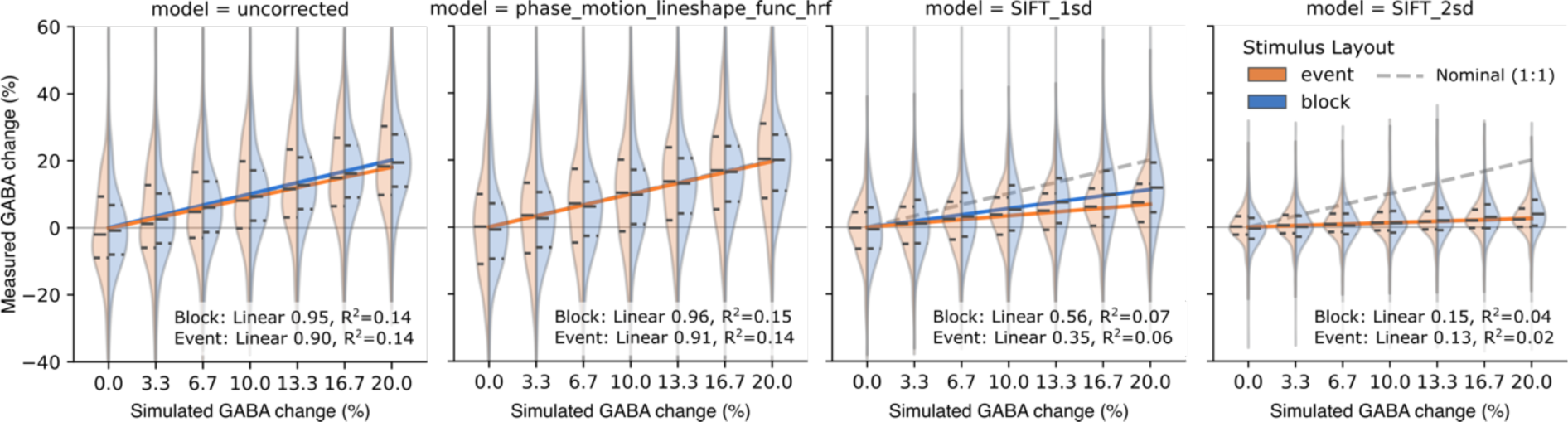
Measured response to simulated GABA change after modelling; linear fit coefficient and coefficient of determination reported as an indication of model sensitivity for simulated block (blue) and event-related (orange) functional stimulus. Response with other models illustrated in Supplementary Figure 4

The SIFT method proved significantly less sensitive to functional changes of interest; with a fairly low 1 SD threshold, only 56% of the simulated functional change was observable (R^2^=0.07) for block stimulus, reduced to 35% (R^2^=0.06) for event-related stimulus. With a more typical 2 SD threshold, sensitivity was further reduced to 15% (R^2^=0.04) for block designs and 13% (R^2^=0.02) for event-related designs.

## 4 Discussion

The present study investigates a linear-modelling approach to characterise variability between transients within an MRS acquisition. Despite this being recognised as an ill-posed inverse problem, the proposed approach is shown to be effective in reducing unwanted variance, whilst retaining the temporal dynamics of interest in fMRS contexts.

### 4.1 Models for reduction of variance

Estimates for within-scan CV before modelling variance are consistent with previous reports^51^. For static MRS measurements, the SIFT approach to removing variance clearly outperformed the proposed linear modelling approach across all metrics, making it the natural choice in such scenarios. However, substantially reduced sensitivity in functional contexts may limit its usability in these applications, and we suggest that this method be applied with particular caution to avoid type-two errors.

Incorporation of phase cycle step alone in the covariate model yielded robust improvements in many scenarios. Since this factor is readily defined without further modelling or inference, its inclusion represents a straightforward addition to improve outcomes over existing pipelines. Nonetheless, the most robust improvements in SNR and reliability were seen in composite models including inferred motion and some means of lineshape matching in addition to the phase cycle factor. We note that the motion covariate alone was not beneficial, and that despite measurable improvements in other quality metrics, the model did not substantially improve the rejection rate in the low-SNR unbalanced edit-ON/edit-OFF case.

### 4.2 Lineshape matching: when and how?

We have previously demonstrated that the choice of line-broadening function for matching will have an effect on concentration estimates, and that an inappropriate choice may introduce bias in quantitative outcomes ^24^. The various matching approaches tested herein differed only marginally in terms of quality of the resultant spectra, and repeatability of the derived concentration estimates. We therefore recommend adoption of the technique which best matches the theoretically anticipated changes. In many situations, linewidth changes are likely to reflect a combination of T2 (Lorentzian), T2* (Gaussian) and other, perhaps asymmetric factors; a generalised matching based on a deconvolution approach is best suited to capturing all these aspects. Post-hoc matching of the modelled spectra (rather than early matching of the partitioned segments) appeared to perform marginally better, likely benefitting from greater SNR in the modelled spectra allowing more robust modelling of the reference peak shape.

### 4.3 Limitations and further work

While the analysis here was limited to GABA-edited (MEGA-PRESS) data, the same techniques may be extended to multi-editing sequences (e.g., HERMES, HERCULES), applied with different localisation schemes (e.g., sLASER), or applied to unedited data. Although similar benefits may be anticipated in many of these cases, further investigation would be required to confirm this – particularly for the unedited case, without the risk of editing artefacts arising from mismatched sub-spectra.

The indirect assessment of “inferred movement” from the estimated centre frequency may have limited accuracy. In principle, a more reliable model for movement could be obtained from motion tracking devices or periodic navigators, and the model verified with subjects moving at defined times (perhaps with a training approach similar to Tapper et al, 2019^52^). However, while the data-driven approach may not perfectly or exclusively model motion, the broader interpretation as “changing scan conditions” remains relevant as a basis for covariate modelling. Regardless, special consideration would be needed for data acquired with real-time frequency adjustment (often presenting as a distinctive sawtooth pattern in the observed water frequency plot).

Although the current analysis does not consider circulatory or respiratory effects, with subtle motion, blood flow and oxygenation potentially impacting signal relaxation rates and spectral lineshape, these and other factors could be trivially incorporated into the covariate model if available. Inclusion of additional factors should be approached with care to avoid overfitting.

The current models operate pointwise on frequency-domain transients, to yield 1D spectra which can then be fit with most existing fitting algorithms. This has the advantage that it may be directly incorporated into existing pipelines, with well-established fitting methods. However, the same factors could in principle be used to inform models for full 2D fitting techniques ^53–56^ which are increasingly being adopted in functional contexts – where explicitly incorporating frequency/phase components into the 2D modelling has been shown to yield improved performance in low SNR cases^57^.

### 4.4 Conclusions

The linear modelling approach presented here is demonstrably effective at improving SNR and test-retest reliability (within scan) of MRS measurements. Moreover, the proposed approach preserves functional changes of interest, outperforming established methods in that regard. We therefore recommend that linear modelling with covariates for intrinsic cyclic effects at a minimum, as well as additional covariates for subject motion if available, should be incorporated into fMRS processing pipelines in preference to simple binning/averaging, where final quantification is to be performed with existing 1D fitting methods.

## List of Abbreviations

(f)MRS (functional): Magnetic Resonance Spectroscopy
BOLD: Blood Oxygenation Level Dependent
CI_95%(,boot)_: 95% confidence interval ((boot) denotes bootstrap CI)
CV: Coefficient of Variation
DIFF: Edited difference spectrum
FID: Free Induction Decay
FWHM_(metab)_: Full Width at Half Maximum linewidth (for the given metabolite)
GABA: γ-aminobutyric acid
GABA+: total edited signal at 3 ppm; GABA with underlying co-edited signals
HRF: Haemodynamic Response Function, characteristic of BOLD response
MAD: Median Absolute Deviation
MSD: Mean Signed Difference
OSF: Rank-Order Statistical Filtering
p_holm_: Holm-Bonferroni adjusted p-value
SIFT: Spectral Improvement by Fourier Thresholding
SNR: signal-to-noise ratio
SNR_(metab)_: Signal-to-Noise Ratio (for given metabolite)

## 5 Acknowledgements

Analysis was performed within a project funded by ERC grant #693124, which additionally supported the contributions of ARC, LE, KH. Data used in this analysis were previously collected through an international collaborative study funded under NIH grant R01 EB016089.

## 6 Declaration of Interest

The authors declare no conflicting interests.

## 7 Data Availability Statement

Scripts used for the present analysis are publicly available here; further dependencies are described within: https://git.app.uib.no/bergen-fmri/gaba-temporal-variability Spectra analysed in this manuscript were obtained from the publicly available Big GABA repository on NITRC, https://www.nitrc.org/projects/biggaba Vendor-specific basis sets used to simulate functional changes were obtained from the publicly available Osprey package, https://schorschinho.github.io/osprey

## A MRSinMRS checklist

**Supplementary Table 1:**
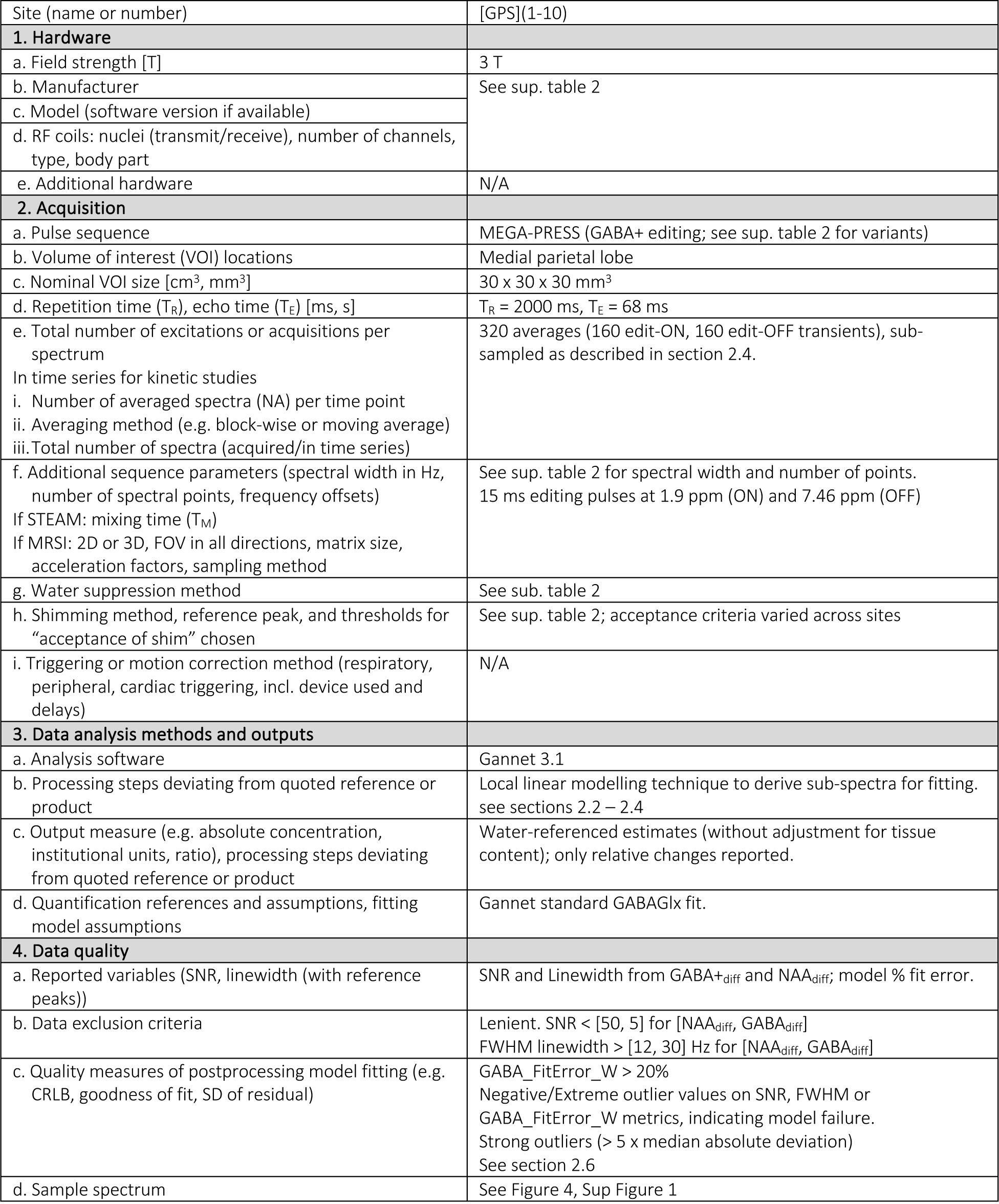
MRSinMRS checklist^18^ summarising key details of the MRS acquisition; reproduced from ^24^.

## B Acquisition Details

**Supplementary Table 2:**
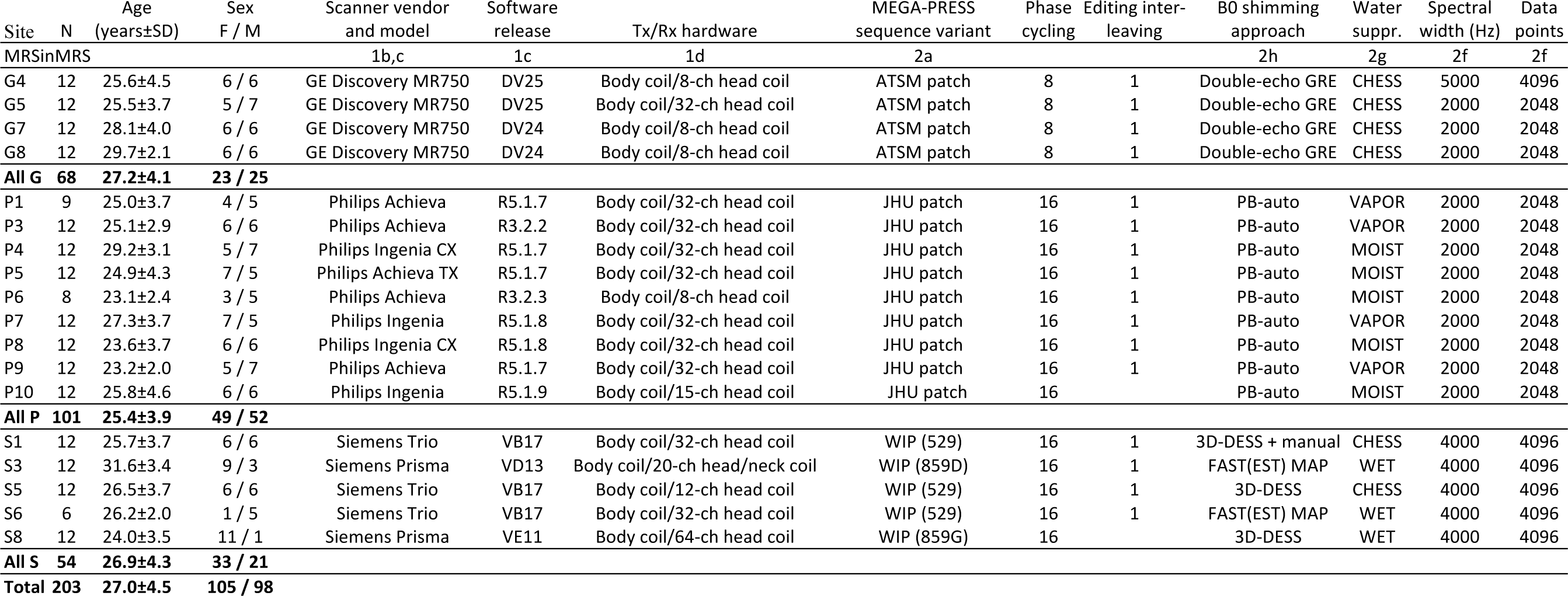
Basic demographics, hardware and software parameters for the constituent datasets; reproduced from previously published work ^17^.

| Site | N | Age<br>(years±SD) | Sex<br>F / M | Scanner vendor<br>and model | Software<br>release | Tx/Rx hardware | MEGA-PRESS<br>sequence variant | Phase<br>cycling | Editing inter-<br>leaving | B0 shimming<br>approach | Water<br>suppr. | Spectral<br>width (Hz) | Data<br>points |
| --- | --- | --- | --- | --- | --- | --- | --- | --- | --- | --- | --- | --- | --- |
| MRSinMRS |  |  |  | 1b,c | 1c | 1d | 2a |  |  | 2h | 2g | 2f | 2f |
| G4 | 12 | 25.6±4.5 | 6 / 6 | GE Discovery MR750 | DV25 | Body coil/8-ch head coil | ATSM patch | 8 | 1 | Double-echo GRE | CHESS | 5000 | 4096 |
| G5 | 12 | 25.5±3.7 | 5 / 7 | GE Discovery MR750 | DV25 | Body coil/32-ch head coil | ATSM patch | 8 | 1 | Double-echo GRE | CHESS | 2000 | 2048 |
| G7 | 12 | 28.1±4.0 | 6 / 6 | GE Discovery MR750 | DV24 | Body coil/8-ch head coil | ATSM patch | 8 | 1 | Double-echo GRE | CHESS | 2000 | 2048 |
| G8 | 12 | 29.7±2.1 | 6 / 6 | GE Discovery MR750 | DV24 | Body coil/8-ch head coil | ATSM patch | 8 | 1 | Double-echo GRE | CHESS | 2000 | 2048 |
| **All G** | **68** | **27.2±4.1** | **23 / 25** |  |  |  |  |  |  |  |  |  |  |
| P1 | 9 | 25.0±3.7 | 4 / 5 | Philips Achieva | R5.1.7 | Body coil/32-ch head coil | JHU patch | 16 | 1 | PB-auto | VAPOR | 2000 | 2048 |
| P3 | 12 | 25.1±2.9 | 6 / 6 | Philips Achieva | R3.2.2 | Body coil/32-ch head coil | JHU patch | 16 | 1 | PB-auto | VAPOR | 2000 | 2048 |
| P4 | 12 | 29.2±3.1 | 5 / 7 | Philips Ingenia CX | R5.1.7 | Body coil/32-ch head coil | JHU patch | 16 | 1 | PB-auto | MOIST | 2000 | 2048 |
| P5 | 12 | 24.9±4.3 | 7 / 5 | Philips Achieva TX | R5.1.7 | Body coil/32-ch head coil | JHU patch | 16 | 1 | PB-auto | MOIST | 2000 | 2048 |
| P6 | 8 | 23.1±2.4 | 3 / 5 | Philips Achieva | R3.2.3 | Body coil/8-ch head coil | JHU patch | 16 | 1 | PB-auto | MOIST | 2000 | 2048 |
| P7 | 12 | 27.3±3.7 | 7 / 5 | Philips Ingenia | R5.1.8 | Body coil/32-ch head coil | JHU patch | 16 | 1 | PB-auto | VAPOR | 2000 | 2048 |
| P8 | 12 | 23.6±3.7 | 6 / 6 | Philips Ingenia CX | R5.1.8 | Body coil/32-ch head coil | JHU patch | 16 | 1 | PB-auto | MOIST | 2000 | 2048 |
| P9 | 12 | 23.2±2.0 | 5 / 7 | Philips Achieva | R5.1.7 | Body coil/32-ch head coil | JHU patch | 16 | 1 | PB-auto | VAPOR | 2000 | 2048 |
| P10 | 12 | 25.8±4.6 | 6 / 6 | Philips Ingenia | R5.1.9 | Body coil/15-ch head coil | JHU patch | 16 |  | PB-auto | MOIST | 2000 | 2048 |
| **All P** | **101** | **25.4±3.9** | **49 / 52** |  |  |  |  |  |  |  |  |  |  |
| S1 | 12 | 25.7±3.7 | 6 / 6 | Siemens Trio | VB17 | Body coil/32-ch head coil | WIP (529) | 16 | 1 | 3D-DESS + manual | CHESS | 4000 | 4096 |
| S3 | 12 | 31.6±3.4 | 9 / 3 | Siemens Prisma | VD13 | Body coil/20-ch head/neck coil | WIP (859D) | 16 | 1 | FAST(EST) MAP | WET | 4000 | 4096 |
| S5 | 12 | 26.5±3.7 | 6 / 6 | Siemens Trio | VB17 | Body coil/12-ch head coil | WIP (529) | 16 | 1 | 3D-DESS | CHESS | 4000 | 4096 |
| S6 | 6 | 26.2±2.0 | 1 / 5 | Siemens Trio | VB17 | Body coil/32-ch head coil | WIP (529) | 16 | 1 | FAST(EST) MAP | WET | 4000 | 4096 |
| S8 | 12 | 24.0±3.5 | 11 / 1 | Siemens Prisma | VE11 | Body coil/64-ch head coil | WIP (859G) | 16 |  | 3D-DESS | WET | 4000 | 4096 |
| **All S** | **54** | **26.9±4.3** | **33 / 21** |  |  |  |  |  |  |  |  |  |  |
| **Total** | **203** | **27.0±4.5** | **105 / 98** |  |  |  |  |  |  |  |  |  |  |

## C Supplementary Results

### C.1 Modelled spectra (single subject)

**Supplementary Figure 1.**
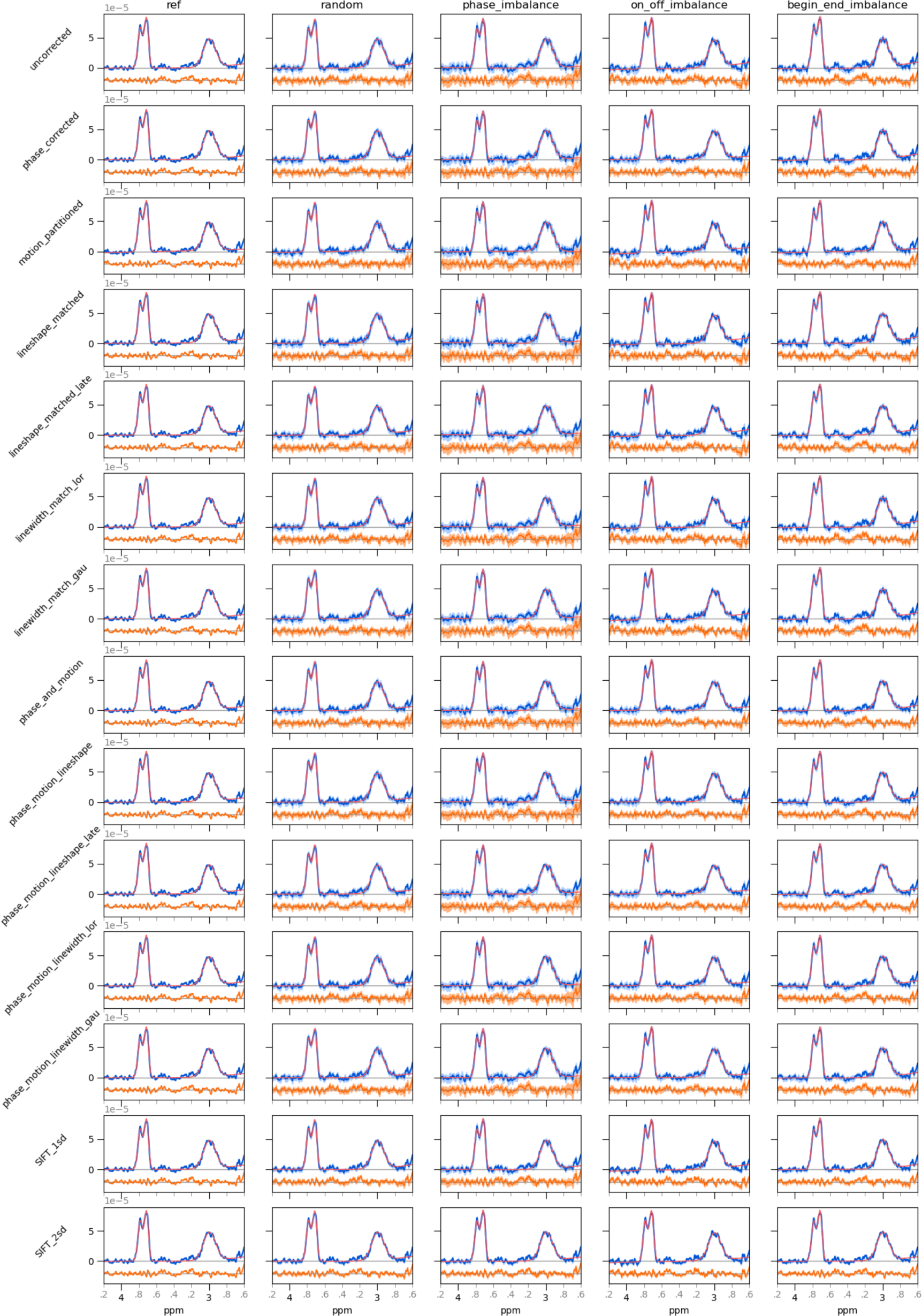
Reconstructed spectra and residuals, for each combination of modelled factors and sampling scenarios for a single representative subject. Mean data (blue) and residuals (orange, lower part) are presented, with standard deviation across repetitions shaded and mean GABAGlx model fit shown in red.

### C.2 Quality of modelled spectra and fits (all subjects)

**Supplementary Table 3.**
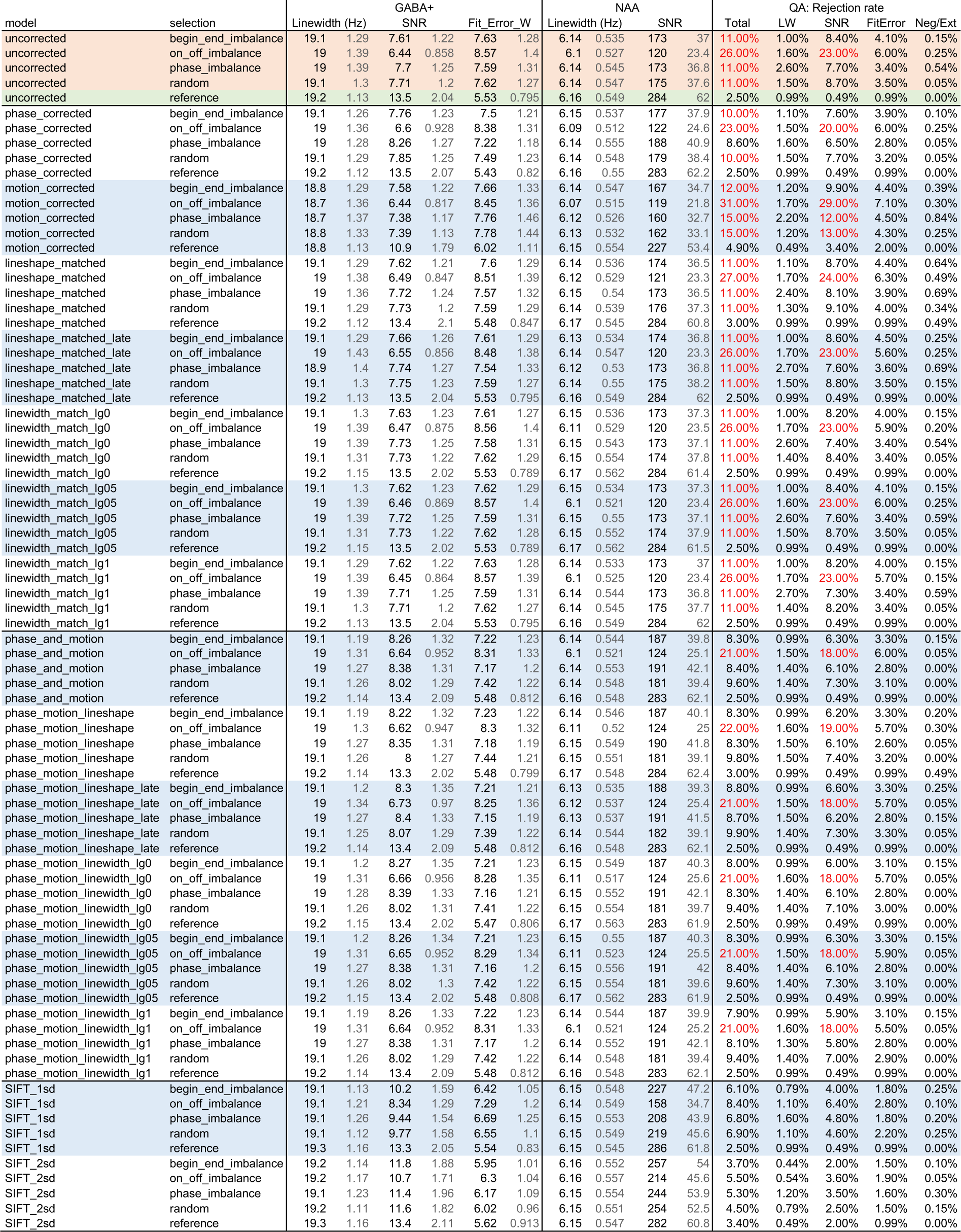
Spectral Quality and rejection rates according to model and sampling scenario; values quoted are median and MAD (lighter font); high rejection rates (>10%) indicated in red text.

### C.3 Model Performance: Signal-to-Noise Ratio

**Supplementary Table 4.** Model performance: Signal-to-Noise Ratio.

| model | selection | uncorrected | filtered | N | Diff % | Diff % 95% CI | punc | Pholm | sig. |
| --- | --- | --- | --- | --- | --- | --- | --- | --- | --- |
| phase_corrected | random | 7.57 ± 1.06 | 7.78 ± 1.16 | 189 | 2.67 | [ 2.22, 3.09] | 3.99E-23 | 1.08E-21 | *** |
| motion_partitioned | random | 7.58 ± 1.07 | 7.23 ± 1.19 | 188 | -5.8 | [-6.80, -4.85] | 1 | 1 |  |
| lineshape_matched | random | 7.54 ± 1.07 | 7.58 ± 1.06 | 188 | 0.21 | [-0.09, 0.47] | 0.00302 | 0.0272 | * |
| lineshape_matched_late | random | 7.57 ± 1.06 | 7.63 ± 1.1 | 189 | 0.49 | [ 0.10, 0.85] | 0.00339 | 0.0272 | * |
| linewidth_match_lor | random | 7.57 ± 1.06 | 7.6 ± 1.06 | 189 | 0.44 | [ 0.25, 0.84] | 0.00000349 | 0.0000559 | *** |
| linewidth_match_gau | random | 7.57 ± 1.06 | 7.57 ± 1.06 | 189 | 0.04 | [ 0.01, 0.12] | 0.00000138 | 0.0000235 | *** |
| linewidth_match_LGmix | random | 7.57 ± 1.06 | 7.59 ± 1.07 | 189 | 0.28 | [ 0.16, 0.57] | 0.00000599 | 0.0000898 | *** |
| phase_and_motion | random | 7.57 ± 1.06 | 8.05 ± 1.2 | 189 | 4.58 | [ 4.10, 5.02] | 4.95E-30 | 2.08E-28 | *** |
| phase_motion_lineshape | random | 7.57 ± 1.06 | 8.04 ± 1.22 | 189 | 4.09 | [ 2.95, 4.64] | 6.69E-28 | 2.37E-26 | *** |
| phase_motion_lineshape_late | random | 7.57 ± 1.06 | 7.94 ± 1.18 | 189 | 5 | [ 4.52, 5.48] | 1.34E-30 | 6.15E-29 | *** |
| phase_motion_linewidth_lor | random | 7.57 ± 1.06 | 8.05 ± 1.2 | 189 | 5.12 | [ 4.60, 5.74] | 1.39E-30 | 6.24E-29 | *** |
| phase_motion_linewidth_gau | random | 7.57 ± 1.06 | 8.05 ± 1.2 | 189 | 4.62 | [ 4.16, 5.08] | 4.95E-30 | 2.08E-28 | *** |
| phase_motion_linewidth_LGmix | random | 7.57 ± 1.06 | 8.05 ± 1.2 | 189 | 4.93 | [ 4.42, 5.49] | 2.02E-30 | 8.88E-29 | *** |
| SIFT_1sd | random | 7.57 ± 1.06 | 9.86 ± 1.48 | 189 | 28.76 | [27.77,29.58] | 4.72E-33 | 2.65E-31 | *** |
| SIFT_2sd | random | 7.57 ± 1.06 | 11.9 ± 1.71 | 189 | 54.58 | [53.01,55.94] | 4.57E-33 | 2.65E-31 | *** |
| phase_corrected | phase_imbalance | 7.52 ± 1.12 | 8.14 ± 1.16 | 190 | 7.93 | [ 6.98, 8.90] | 9.05E-28 | 3.08E-26 | *** |
| motion_partitioned | phase_imbalance | 7.58 ± 1.13 | 7.27 ± 1.13 | 188 | -5.78 | [-6.83, -4.82] | 1 | 1 |  |
| lineshape_matched | phase_imbalance | 7.48 ± 1.13 | 7.59 ± 1.13 | 189 | 0.2 | [-0.14, 0.52] | 0.00453 | 0.0317 | * |
| lineshape_matched_late | phase_imbalance | 7.52 ± 1.12 | 7.6 ± 1.17 | 190 | 0.62 | [ 0.27, 0.96] | 0.0000277 | 0.000305 | *** |
| linewidth_match_lor | phase_imbalance | 7.52 ± 1.12 | 7.58 ± 1.13 | 190 | 0.5 | [ 0.27, 0.90] | 0.0000125 | 0.000163 | *** |
| linewidth_match_gau | phase_imbalance | 7.52 ± 1.12 | 7.52 ± 1.12 | 190 | 0.04 | [ 0.01, 0.09] | 0.00000666 | 0.0000932 | *** |
| linewidth_match_LGmix | phase_imbalance | 7.52 ± 1.12 | 7.55 ± 1.13 | 190 | 0.3 | [ 0.15, 0.56] | 0.0000235 | 0.000282 | *** |
| phase_and_motion | phase_imbalance | 7.52 ± 1.12 | 8.26 ± 1.17 | 190 | 9.31 | [ 8.32,10.30] | 2.6E-29 | 9.63E-28 | *** |
| phase_motion_lineshape | phase_imbalance | 7.52 ± 1.12 | 8.24 ± 1.17 | 190 | 8.78 | [ 7.46, 9.81] | 6.59E-28 | 2.37E-26 | *** |
| phase_motion_lineshape_late | phase_imbalance | 7.52 ± 1.12 | 8.32 ± 1.24 | 190 | 10.1 | [ 9.08,11.13] | 4.85E-30 | 2.08E-28 | *** |
| phase_motion_linewidth_lor | phase_imbalance | 7.52 ± 1.12 | 8.26 ± 1.19 | 190 | 9.87 | [ 8.85,10.95] | 6.6E-30 | 2.64E-28 | *** |
| phase_motion_linewidth_gau | phase_imbalance | 7.52 ± 1.12 | 8.26 ± 1.16 | 190 | 9.36 | [ 8.36,10.36] | 1.91E-29 | 7.24E-28 | *** |
| phase_motion_linewidth_LGmix | phase_imbalance | 7.52 ± 1.12 | 8.26 ± 1.15 | 190 | 9.69 | [ 8.68,10.70] | 8.26E-30 | 3.22E-28 | *** |
| SIFT_1sd | phase_imbalance | 7.52 ± 1.12 | 9.33 ± 1.48 | 190 | 23.46 | [22.56,24.33] | 3.18E-33 | 1.88E-31 | *** |
| SIFT_2sd | phase_imbalance | 7.52 ± 1.12 | 11.5 ± 1.87 | 190 | 49.63 | [47.81,51.25] | 3.13E-33 | 1.88E-31 | *** |
| phase_corrected | on_off_imbalance | 6.19 ± 0.79 | 6.33 ± 0.779 | 181 | 2.57 | [ 2.10, 2.99] | 5.99E-21 | 1.5E-19 | *** |
| motion_partitioned | on_off_imbalance | 6.24 ± 0.756 | 6.2 ± 0.733 | 174 | -1.92 | [-2.82, -1.12] | 1 | 1 |  |
| lineshape_matched | on_off_imbalance | 6.2 ± 0.769 | 6.26 ± 0.727 | 177 | 0.39 | [ 0.08, 0.68] | 0.000206 | 0.00206 | ** |
| lineshape_matched_late | on_off_imbalance | 6.19 ± 0.79 | 6.38 ± 0.739 | 181 | 1.77 | [ 1.33, 2.20] | 3.37E-14 | 8.08E-13 | *** |
| linewidth_match_lor | on_off_imbalance | 6.19 ± 0.79 | 6.21 ± 0.801 | 181 | 0.71 | [ 0.47, 1.08] | 0.00000001 | 0.00000041 | *** |
| linewidth_match_gau | on_off_imbalance | 6.19 ± 0.79 | 6.2 ± 0.794 | 181 | 0.07 | [ 0.04, 0.15] | 0.00000003 | 0.00000082 | *** |
| linewidth_match_LGmix | on_off_imbalance | 6.19 ± 0.79 | 6.2 ± 0.81 | 181 | 0.44 | [ 0.29, 0.70] | 0.00000008 | 0.00000175 | *** |
| phase_and_motion | on_off_imbalance | 6.19 ± 0.784 | 6.4 ± 0.806 | 180 | 4.17 | [ 3.61, 4.67] | 3.93E-25 | 1.14E-23 | *** |
| phase_motion_lineshape | on_off_imbalance | 6.19 ± 0.78 | 6.42 ± 0.784 | 179 | 3.65 | [ 2.48, 4.27] | 5.24E-23 | 1.36E-21 | *** |
| phase_motion_lineshape_late | on_off_imbalance | 6.19 ± 0.784 | 6.49 ± 0.773 | 180 | 5.26 | [ 4.64, 5.83] | 6.2E-26 | 1.92E-24 | *** |
| phase_motion_linewidth_lor | on_off_imbalance | 6.19 ± 0.784 | 6.42 ± 0.825 | 180 | 4.74 | [ 4.14, 5.40] | 2.55E-27 | 8.43E-26 | *** |
| phase_motion_linewidth_gau | on_off_imbalance | 6.19 ± 0.784 | 6.4 ± 0.806 | 180 | 4.2 | [ 3.65, 4.74] | 2.65E-25 | 7.96E-24 | *** |
| phase_motion_linewidth_LGmix | on_off_imbalance | 6.19 ± 0.784 | 6.4 ± 0.814 | 180 | 4.55 | [ 3.98, 5.16] | 9.33E-27 | 2.98E-25 | *** |
| SIFT_1sd | on_off_imbalance | 6.19 ± 0.784 | 8.42 ± 1.17 | 180 | 34.31 | [32.99,35.48] | 1.37E-31 | 6.7E-30 | *** |
| SIFT_2sd | on_off_imbalance | 6.19 ± 0.784 | 10.9 ± 1.58 | 180 | 73.51 | [71.40,75.36] | 1.37E-31 | 6.7E-30 | *** |
| phase_corrected | begin_end_imbalance | 7.5 ± 1.13 | 7.74 ± 1.16 | 190 | 2.47 | [ 2.11, 2.84] | 2.4E-23 | 6.71E-22 | *** |
| motion_partitioned | begin_end_imbalance | 7.5 ± 1.13 | 7.52 ± 1.22 | 188 | -2.39 | [-3.60, -1.25] | 1 | 1 |  |
| lineshape_matched | begin_end_imbalance | 7.5 ± 1.13 | 7.54 ± 1.11 | 189 | -0.11 | [-0.61, 0.24] | 0.125 | 0.625 |  |
| lineshape_matched_late | begin_end_imbalance | 7.5 ± 1.13 | 7.51 ± 1.11 | 190 | 0.42 | [ 0.06, 0.86] | 0.0619 | 0.372 |  |
| linewidth_match_lor | begin_end_imbalance | 7.5 ± 1.13 | 7.55 ± 1.1 | 190 | 0.53 | [ 0.30, 0.95] | 0.00000027 | 0.00000556 | *** |
| linewidth_match_gau | begin_end_imbalance | 7.5 ± 1.13 | 7.5 ± 1.13 | 190 | 0.07 | [ 0.02, 0.18] | 0.00000049 | 0.00000932 | *** |
| linewidth_match_LGmix | begin_end_imbalance | 7.5 ± 1.13 | 7.52 ± 1.11 | 190 | 0.34 | [ 0.20, 0.60] | 0.00000070 | 0.0000127 | *** |
| phase_and_motion | begin_end_imbalance | 7.5 ± 1.13 | 8.15 ± 1.16 | 189 | 9.27 | [ 8.57, 9.95] | 1.47E-32 | 7.51E-31 | *** |
| phase_motion_lineshape | begin_end_imbalance | 7.5 ± 1.13 | 8.11 ± 1.13 | 189 | 8.68 | [ 7.48, 9.48] | 5.12E-31 | 2.41E-29 | *** |
| phase_motion_lineshape_late | begin_end_imbalance | 7.5 ± 1.13 | 8.17 ± 1.17 | 189 | 9.7 | [ 9.00,10.41] | 1.08E-32 | 5.82E-31 | *** |
| phase_motion_linewidth_lor | begin_end_imbalance | 7.5 ± 1.13 | 8.16 ± 1.17 | 189 | 9.81 | [ 9.05,10.61] | 1.38E-32 | 7.3E-31 | *** |
| phase_motion_linewidth_gau | begin_end_imbalance | 7.5 ± 1.13 | 8.15 ± 1.16 | 189 | 9.35 | [ 8.65,10.05] | 1.44E-32 | 7.51E-31 | *** |
| phase_motion_linewidth_LGmix | begin_end_imbalance | 7.5 ± 1.13 | 8.15 ± 1.17 | 189 | 9.62 | [ 8.91,10.39] | 1.51E-32 | 7.57E-31 | *** |
| SIFT_1sd | begin_end_imbalance | 7.5 ± 1.13 | 10.1 ± 1.52 | 189 | 34.97 | [33.99,35.92] | 4.57E-33 | 2.65E-31 | *** |
| SIFT_2sd | begin_end_imbalance | 7.5 ± 1.13 | 11.9 ± 1.7 | 189 | 58.84 | [57.17,60.38] | 4.57E-33 | 2.65E-31 | *** |

### C.4 Model Performance: Linewidth

**Supplementary Table 5.**
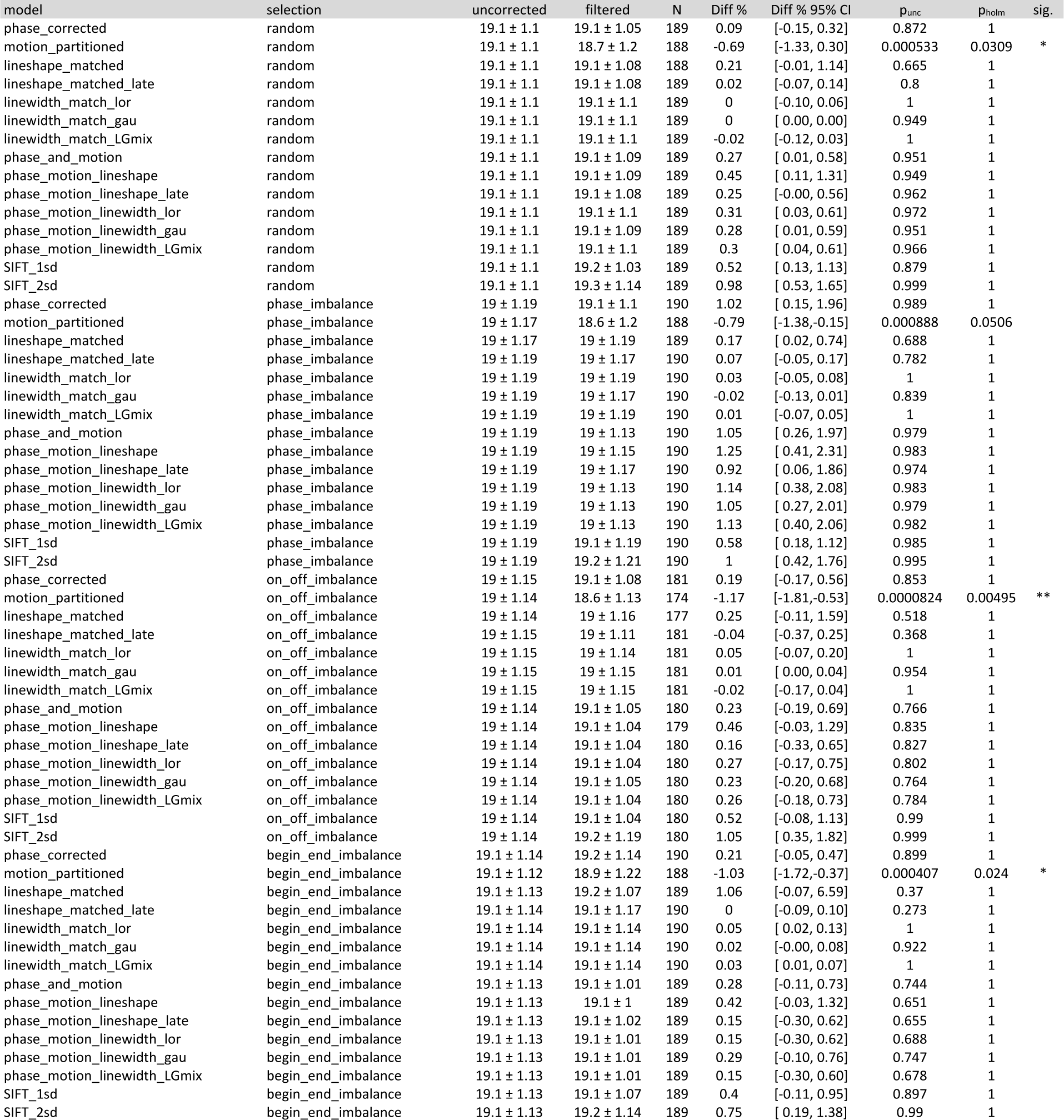
Model performance: FWHM linewidth (Hz)

**Supplementary Figure 2.**
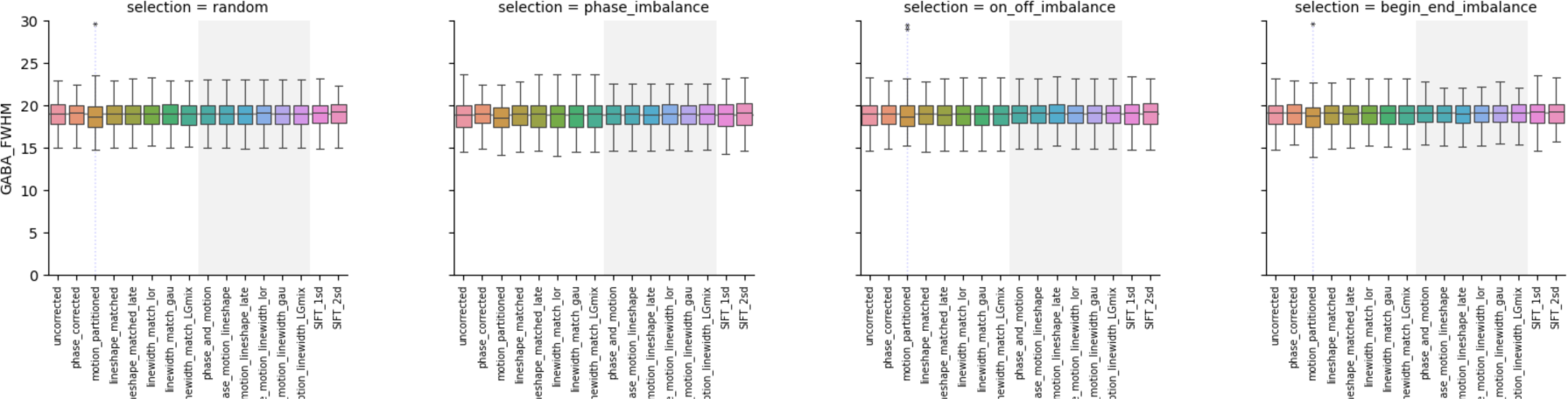
Model performance: FWHM linewidth (Hz)

### C.5 Model Performance: Fit Error

**Supplementary Table 6.**
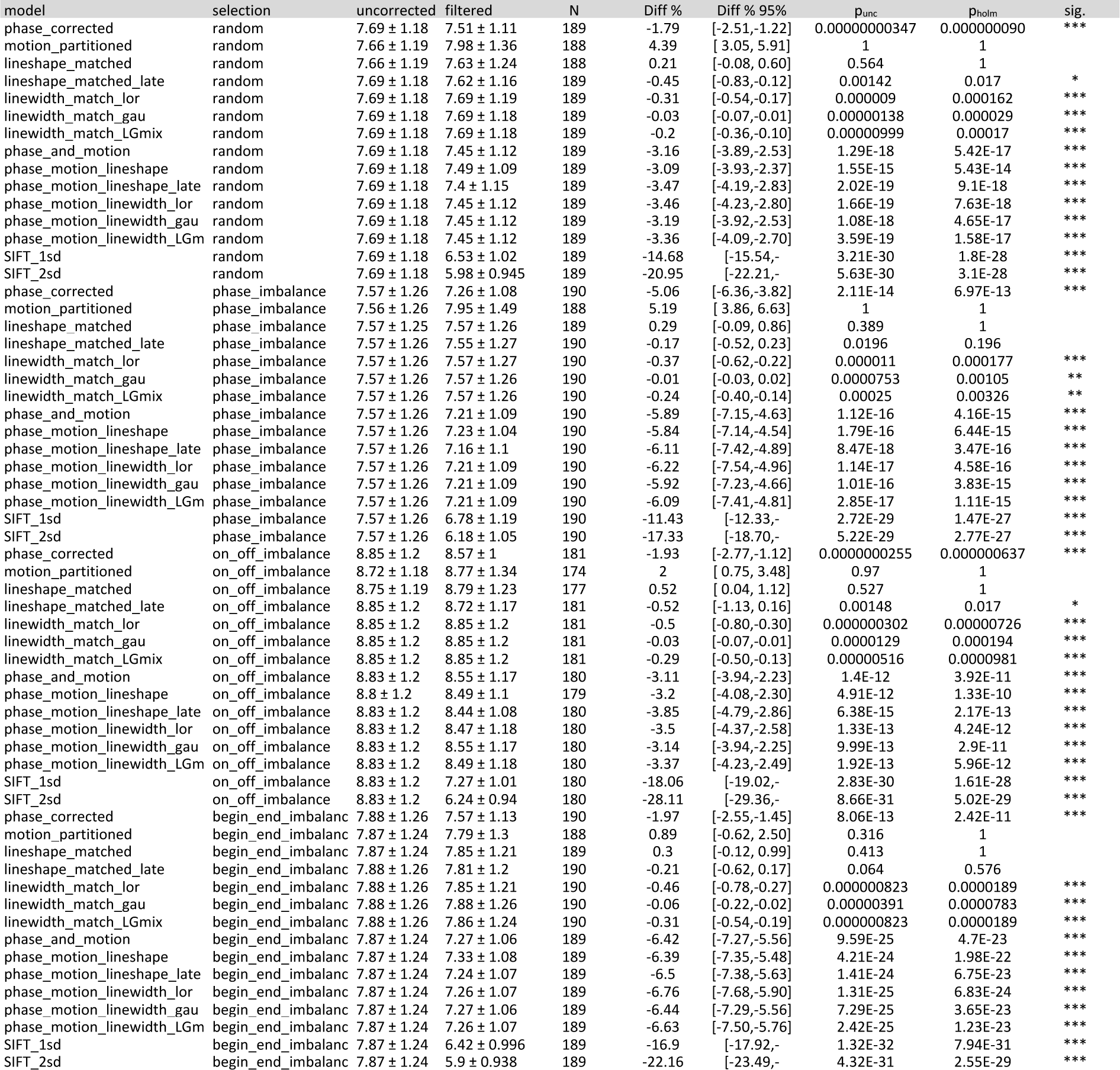
Model performance: GABA Fit Error.

**Supplementary Figure 3.**
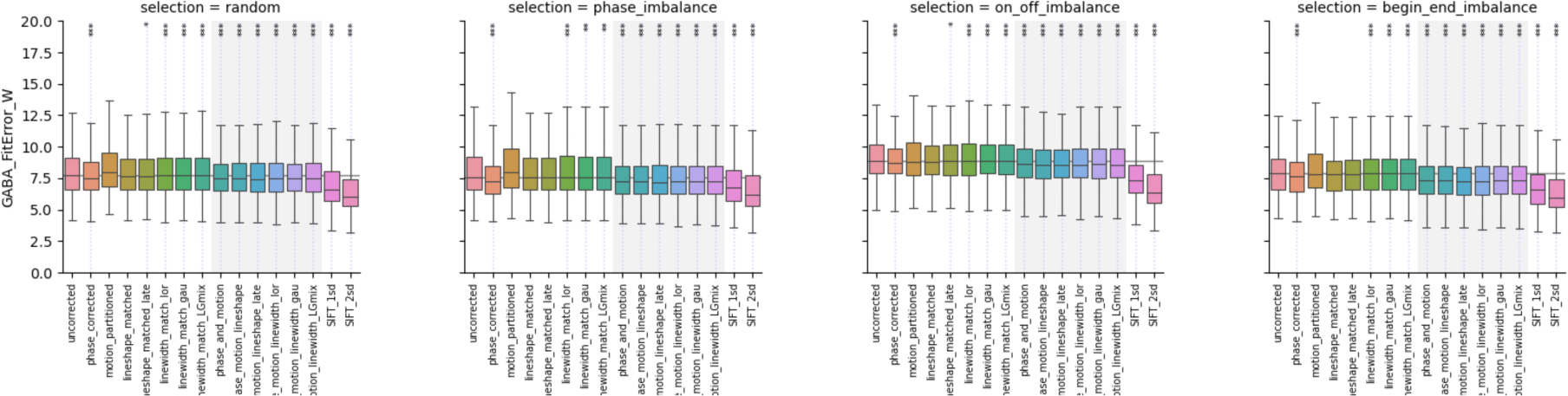
GABA Fit Error.

### C.6 Model Performance: Coefficient of Variation

**Supplementary Table 7.**
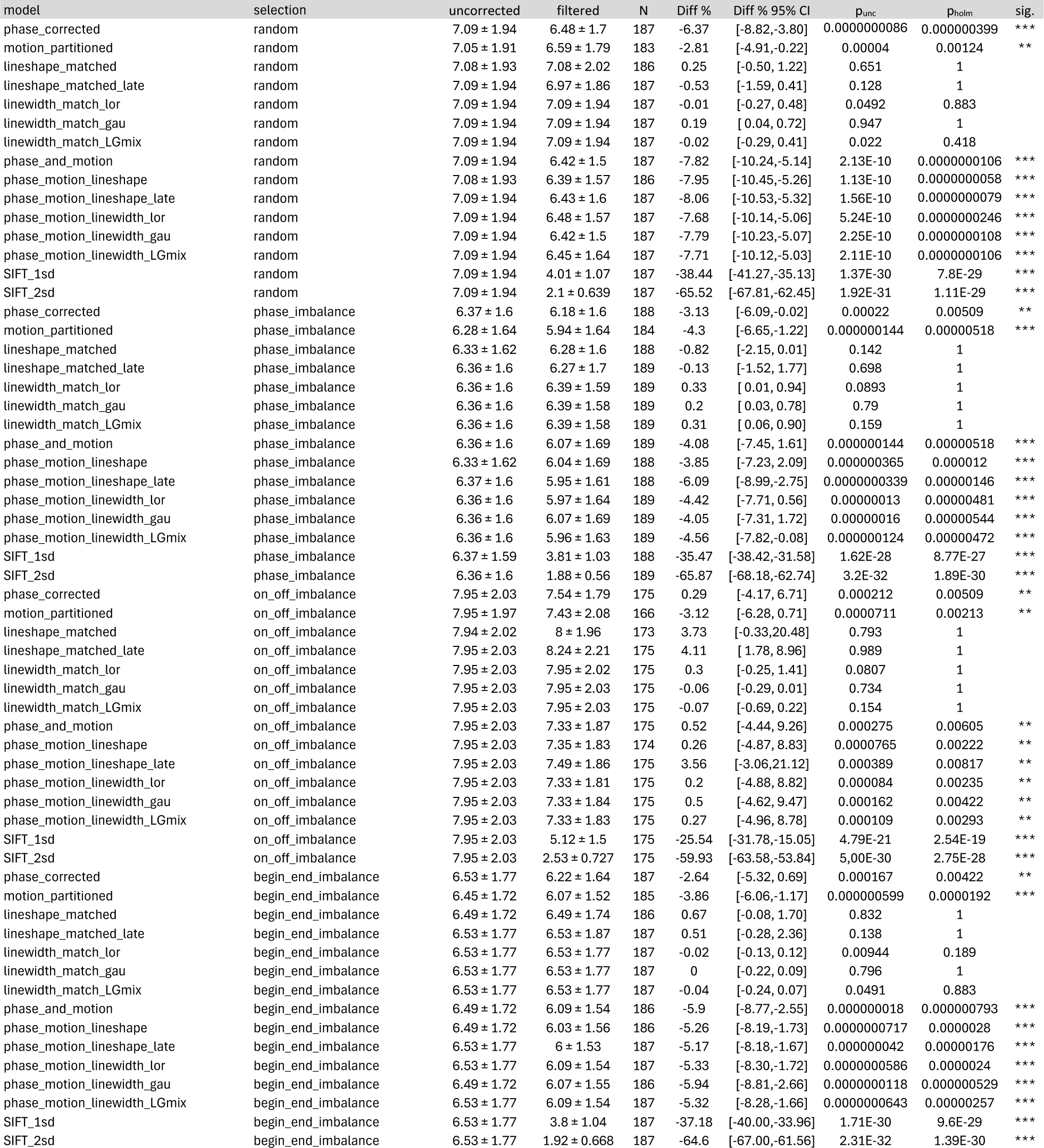
Model performance: Coefficient of Variation.

### C.7 Model Performance: Mean Signed Deviation

**Supplementary Table 8.** Mean Signed Deviation statistics.

| model | selection | filtered | N | MSD % | MSD % 95% CI | punc | Phol | sig. |
| --- | --- | --- | --- | --- | --- | --- | --- | --- |
| phase_corrected | random | -0.136 ± 1.56 | 186 | -0.21 | [-0.73, 0.27] | 0.366 | 1 |  |
| motion_partitioned | random | -0.117 ± 1.51 | 182 | -0.16 | [-0.67, 0.33] | 0.481 | 1 |  |
| lineshape_matched | random | -0.141 ± 1.57 | 185 | -0.21 | [-0.72, 0.29] | 0.363 | 1 |  |
| lineshape_matched_late | random | -0.136 ± 1.56 | 186 | -0.21 | [-0.74, 0.25] | 0.366 | 1 |  |
| linewidth_match_lor | random | -0.136 ± 1.56 | 186 | -0.21 | [-0.72, 0.28] | 0.366 | 1 |  |
| linewidth_match_gau | random | -0.136 ± 1.56 | 186 | -0.21 | [-0.71, 0.27] | 0.366 | 1 |  |
| linewidth_match_LGmix | random | -0.136 ± 1.56 | 186 | -0.21 | [-0.73, 0.27] | 0.366 | 1 |  |
| phase_and_motion | random | -0.136 ± 1.56 | 186 | -0.21 | [-0.73, 0.26] | 0.366 | 1 |  |
| phase_motion_lineshape | random | -0.141 ± 1.57 | 185 | -0.21 | [-0.72, 0.27] | 0.363 | 1 |  |
| phase_motion_lineshape_late | random | -0.136 ± 1.56 | 186 | -0.21 | [-0.73, 0.26] | 0.366 | 1 |  |
| phase_motion_linewidth_lor | random | -0.136 ± 1.56 | 186 | -0.21 | [-0.71, 0.27] | 0.366 | 1 |  |
| phase_motion_linewidth_gau | random | -0.136 ± 1.56 | 186 | -0.21 | [-0.70, 0.29] | 0.366 | 1 |  |
| phase_motion_linewidth_LGmix | random | -0.136 ± 1.56 | 186 | -0.21 | [-0.72, 0.28] | 0.366 | 1 |  |
| SIFT_1sd | random | -0.136 ± 1.56 | 186 | -0.21 | [-0.73, 0.26] | 0.366 | 1 |  |
| SIFT_2sd | random | -0.136 ± 1.56 | 186 | -0.21 | [-0.72, 0.27] | 0.366 | 1 |  |
| phase_corrected | phase_imbalance | 0.226 ± 3.16 | 187 | 0.15 | [-0.84, 1.24] | 0.626 | 1 |  |
| motion_partitioned | phase_imbalance | 0.226 ± 3.09 | 183 | 0.4 | [-0.66, 1.55] | 0.483 | 1 |  |
| lineshape_matched | phase_imbalance | 0.226 ± 3.16 | 187 | 0.44 | [-0.62, 1.59] | 0.466 | 1 |  |
| lineshape_matched_late | phase_imbalance | 0.226 ± 3.16 | 188 | 0.37 | [-0.65, 1.55] | 0.543 | 1 |  |
| linewidth_match_lor | phase_imbalance | 0.226 ± 3.16 | 188 | 0.37 | [-0.70, 1.54] | 0.543 | 1 |  |
| linewidth_match_gau | phase_imbalance | 0.226 ± 3.16 | 188 | 0.37 | [-0.67, 1.53] | 0.543 | 1 |  |
| linewidth_match_LGmix | phase_imbalance | 0.226 ± 3.16 | 188 | 0.37 | [-0.73, 1.50] | 0.543 | 1 |  |
| phase_and_motion | phase_imbalance | 0.226 ± 3.16 | 188 | 0.37 | [-0.69, 1.56] | 0.543 | 1 |  |
| phase_motion_lineshape | phase_imbalance | 0.226 ± 3.16 | 187 | 0.44 | [-0.61, 1.61] | 0.466 | 1 |  |
| phase_motion_lineshape_late | phase_imbalance | 0.226 ± 3.16 | 187 | 0.15 | [-0.86, 1.21] | 0.626 | 1 |  |
| phase_motion_linewidth_lor | phase_imbalance | 0.226 ± 3.16 | 188 | 0.37 | [-0.71, 1.57] | 0.543 | 1 |  |
| phase_motion_linewidth_gau | phase_imbalance | 0.226 ± 3.16 | 188 | 0.37 | [-0.70, 1.57] | 0.543 | 1 |  |
| phase_motion_linewidth_LGmix | phase_imbalance | 0.226 ± 3.16 | 188 | 0.37 | [-0.68, 1.56] | 0.543 | 1 |  |
| SIFT_1sd | phase_imbalance | 0.226 ± 3.16 | 187 | 0.38 | [-0.70, 1.54] | 0.503 | 1 |  |
| SIFT_2sd | phase_imbalance | 0.226 ± 3.16 | 188 | 0.37 | [-0.68, 1.54] | 0.543 | 1 |  |
| phase_corrected | on_off_imbalance | -0.105 ± 3.07 | 174 | 0.38 | [-0.37, 1.12] | 0.284 | 1 |  |
| motion_partitioned | on_off_imbalance | 0.103 ± 2.99 | 165 | 0.55 | [-0.21, 1.30] | 0.179 | 1 |  |
| lineshape_matched | on_off_imbalance | 0.00511 ± 3 | 172 | 0.5 | [-0.21, 1.24] | 0.204 | 1 |  |
| lineshape_matched_late | on_off_imbalance | -0.105 ± 3.07 | 174 | 0.38 | [-0.40, 1.12] | 0.284 | 1 |  |
| linewidth_match_lor | on_off_imbalance | -0.105 ± 3.07 | 174 | 0.38 | [-0.35, 1.14] | 0.284 | 1 |  |
| linewidth_match_gau | on_off_imbalance | -0.105 ± 3.07 | 174 | 0.38 | [-0.40, 1.14] | 0.284 | 1 |  |
| linewidth_match_LGmix | on_off_imbalance | -0.105 ± 3.07 | 174 | 0.38 | [-0.38, 1.13] | 0.284 | 1 |  |
| phase_and_motion | on_off_imbalance | -0.105 ± 3.07 | 174 | 0.38 | [-0.38, 1.13] | 0.284 | 1 |  |
| phase_motion_lineshape | on_off_imbalance | -0.0927 ± 3.08 | 173 | 0.4 | [-0.37, 1.14] | 0.258 | 1 |  |
| phase_motion_lineshape_late | on_off_imbalance | -0.105 ± 3.07 | 174 | 0.38 | [-0.37, 1.13] | 0.284 | 1 |  |
| phase_motion_linewidth_lor | on_off_imbalance | -0.105 ± 3.07 | 174 | 0.38 | [-0.38, 1.13] | 0.284 | 1 |  |
| phase_motion_linewidth_gau | on_off_imbalance | -0.105 ± 3.07 | 174 | 0.38 | [-0.36, 1.11] | 0.284 | 1 |  |
| phase_motion_linewidth_LGmix | on_off_imbalance | -0.105 ± 3.07 | 174 | 0.38 | [-0.37, 1.14] | 0.284 | 1 |  |
| SIFT_1sd | on_off_imbalance | -0.105 ± 3.07 | 174 | 0.38 | [-0.40, 1.13] | 0.284 | 1 |  |
| SIFT_2sd | on_off_imbalance | -0.105 ± 3.07 | 174 | 0.38 | [-0.37, 1.14] | 0.284 | 1 |  |
| phase_corrected | begin_end_imbalance | -0.0829 ± 2.62 | 186 | 0.31 | [-0.33, 1.13] | 0.832 | 1 |  |
| motion_partitioned | begin_end_imbalance | -0.0829 ± 2.62 | 184 | 0.32 | [-0.31, 1.17] | 0.806 | 1 |  |
| lineshape_matched | begin_end_imbalance | -0.0285 ± 2.57 | 185 | 0.34 | [-0.29, 1.19] | 0.751 | 1 |  |
| lineshape_matched_late | begin_end_imbalance | -0.0829 ± 2.62 | 186 | 0.31 | [-0.34, 1.15] | 0.832 | 1 |  |
| linewidth_match_lor | begin_end_imbalance | -0.0829 ± 2.62 | 186 | 0.31 | [-0.34, 1.13] | 0.832 | 1 |  |
| linewidth_match_gau | begin_end_imbalance | -0.0829 ± 2.62 | 186 | 0.31 | [-0.33, 1.15] | 0.832 | 1 |  |
| linewidth_match_LGmix | begin_end_imbalance | -0.0829 ± 2.62 | 186 | 0.31 | [-0.34, 1.12] | 0.832 | 1 |  |
| phase_and_motion | begin_end_imbalance | -0.137 ± 2.6 | 185 | 0.1 | [-0.50, 0.70] | 0.932 | 1 |  |
| phase_motion_lineshape | begin_end_imbalance | -0.0285 ± 2.57 | 185 | 0.34 | [-0.31, 1.16] | 0.751 | 1 |  |
| phase_motion_lineshape_late | begin_end_imbalance | -0.0829 ± 2.62 | 186 | 0.31 | [-0.33, 1.12] | 0.832 | 1 |  |
| phase_motion_linewidth_lor | begin_end_imbalance | -0.0829 ± 2.62 | 186 | 0.31 | [-0.32, 1.14] | 0.832 | 1 |  |
| phase_motion_linewidth_gau | begin_end_imbalance | -0.137 ± 2.6 | 185 | 0.1 | [-0.51, 0.71] | 0.932 | 1 |  |
| phase_motion_linewidth_LGmix | begin_end_imbalance | -0.0829 ± 2.62 | 186 | 0.31 | [-0.34, 1.13] | 0.832 | 1 |  |
| SIFT_1sd | begin_end_imbalance | -0.0829 ± 2.62 | 186 | 0.31 | [-0.33, 1.16] | 0.832 | 1 |  |
| SIFT_2sd | begin_end_imbalance | -0.0829 ± 2.62 | 186 | 0.31 | [-0.34, 1.14] | 0.832 | 1 |  |

### C.8 Fits to simulated functional changes

**Supplementary Table 9.**
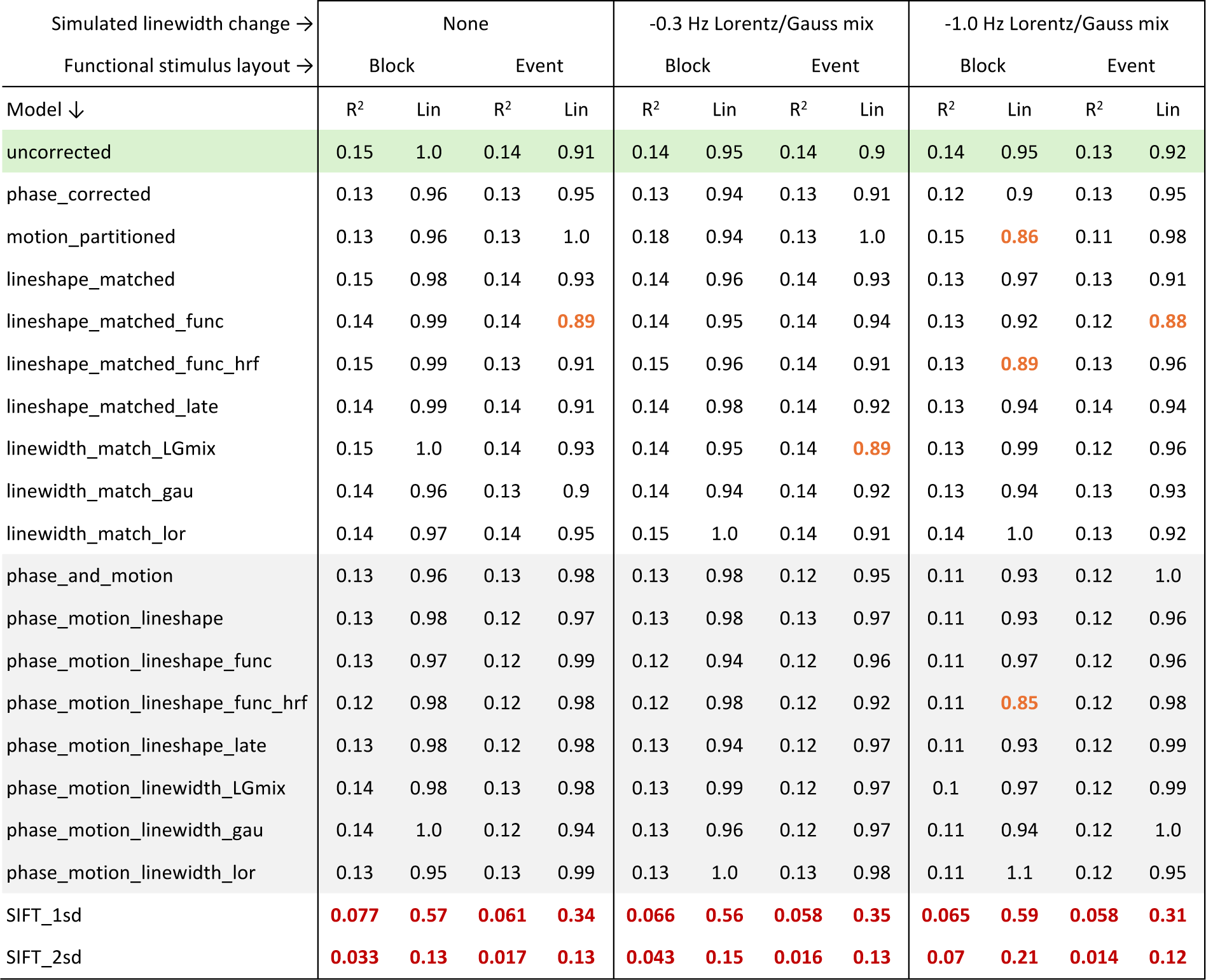
Fits to simulated functional (task-related) GABA+ changes, for the various covariate models. GABA+ changes simulated in response to block or event-related stimulus distribution, with varying degrees of simulated line-broadening as detailed in section 2.5. R^2^ and linear fit coefficient are given after robust regression; sensitivity < 0.9 indicated in orange, sensitivity < 0.8 or R^2^ < 0.1 indicated in red.

**Supplementary Figure 4.**
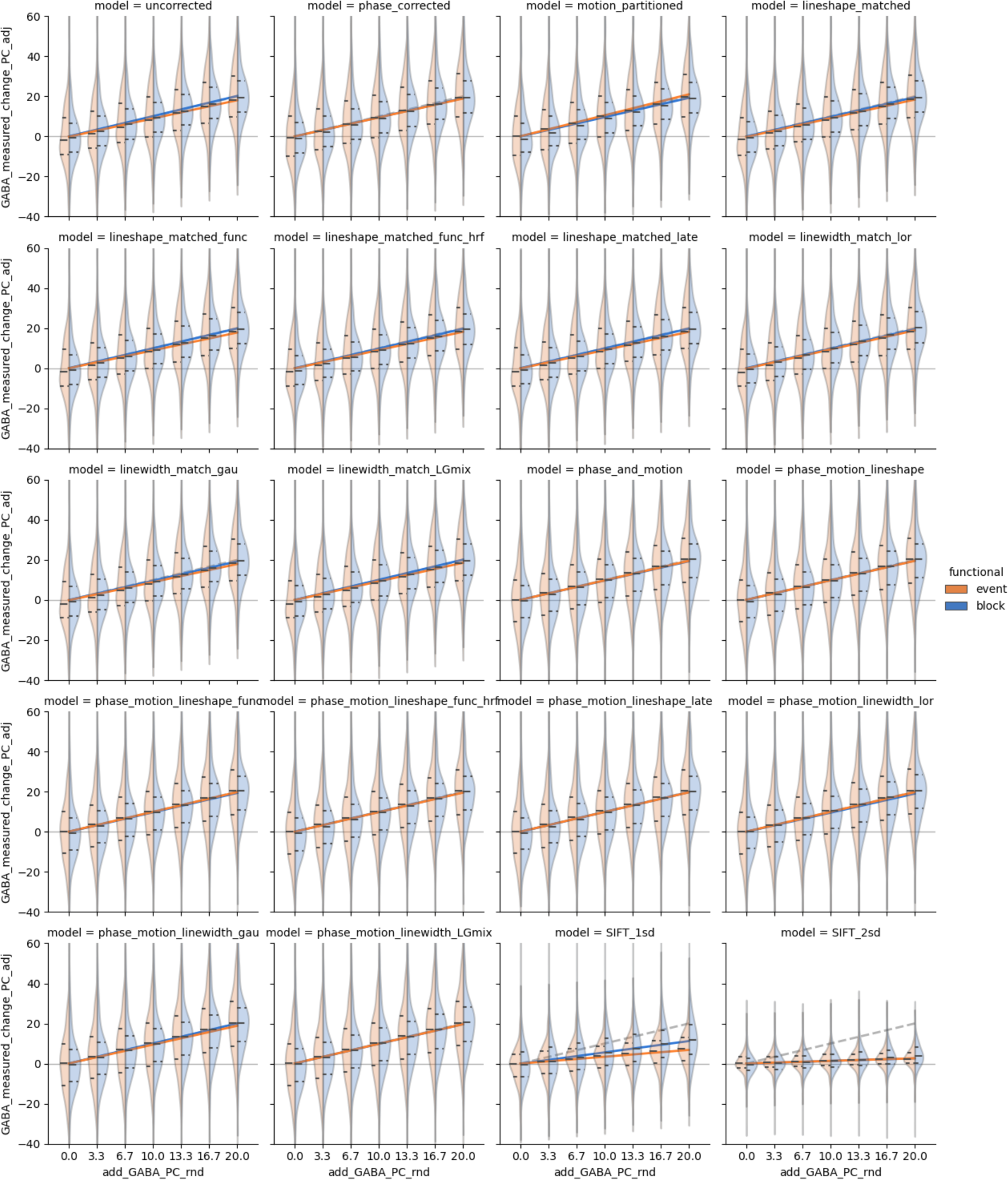
Measured response to simulated GABA change after modelling; corresponding fit coefficients are presented in Supplementary Table 9.

## Notes

### Competing Interest Statement

The authors have declared no competing interest.

## References

1. Hennig J. The application of phase rotation for localized in Vivo proton spectroscopy with short echo times. Journal of Magnetic Resonance (1969). 1992;96(1):40-49. doi:10.1016/0022-2364(92)90286-G

2. Kingsley PB. Product operators, coherence pathways, and phase cycling part i: product operators, spin-spin coupling, and coherence pathways. Concepts Magn Reson. 1995;7(1):29–47. doi:10.1002/cmr.1820070103

3. Kingsley PB. Product operators, coherence pathways, and phase cycling. Part III: phase cycling. Concepts Magn Reson. 1995;7(3):167–192. doi:10.1002/cmr.1820070302

4. Levitt MH. Spin Dynamics: Basics of Nuclear Magnetic Resonance. John Wiley & Sons; 2001.

5. Ramadan S. Phase-rotation in in-vivo localized spectroscopy. Concepts Magn Reson. 2007;30A(3):147-153. doi:10.1002/cmr.a.20080

6. Knight-Scott J, Shanbhag DD, Dunham SA. A phase rotation scheme for achieving very short echo times with localized stimulated echo spectroscopy. Magnetic Resonance Imaging. 2005;23(8):871–876. doi:10.1016/j.mri.2005.07.004

7. Rothman DL, Petroff OA, Behar KL, Mattson RH. Localized 1H NMR measurements of gamma-aminobutyric acid in human brain in vivo. Proceedings of the National Academy of Sciences. 1993;90(12):5662–5666. doi:10.1073/pnas.90.12.5662

8. Mescher M, Merkle H, Kirsch J, Garwood M, Gruetter R. Simultaneous in vivo spectral editing and water suppression. NMR Biomed. 1998;11(6):266–272. doi:10.1002/(sici)1099-1492(199810)11:6<266::aid-nbm530>3.0.co;2-j

9. Apšvalka D, Gadie A, Clemence M, Mullins PG. Event-related dynamics of glutamate and BOLD effects measured using functional magnetic resonance spectroscopy (fMRS) at 3 T in a repetition suppression paradigm. NeuroImage. 2015;118:292–300. doi:10.1016/j.neuroimage.2015.06.015

10. Lally N, Mullins PG, Roberts MV, Price D, Gruber T, Haenschel C. Glutamatergic correlates of gamma-band oscillatory activity during cognition: A concurrent ER-MRS and EEG study. NeuroImage. 2014;85:823–833. doi:10.1016/j.neuroimage.2013.07.049

11. Hui SCN, Mikkelsen M, Zöllner HJ, et al. Frequency drift in MR spectroscopy at 3T. NeuroImage. 2021;241:118430. doi:10.1016/j.neuroimage.2021.118430

12. Slotboom J, Nirkko A, Brekenfeld C, van Ormondt D. Reliability testing of *in vivo* magnetic resonance spectroscopy (MRS) signals and signal artifact reduction by order statistic filtering. Meas Sci Technol. 2009;20(10):104030. doi:10.1088/0957-0233/20/10/104030

13. Doyle M, Chapman BLW, Balschi JA, Pohost GM. SIFT, a Postprocessing Method That Increases the Signal-to-Noise Ratio of Spectra Which Vary in Time. Journal of Magnetic Resonance, Series B. 1994;103(2):128–133. doi:10.1006/jmrb.1994.1020

14. Rowland B, Merugumala SK, Liao H, Creager MA, Balschi J, Lin AP. Spectral improvement by fourier thresholding of in vivo dynamic spectroscopy data: SIFT Denoising for Dynamic MRS Data. Magn Reson Med. 2016;76(3):978–985. doi:10.1002/mrm.25976

15. Mikkelsen M, Barker PB, Bhattacharyya PK, et al. Big GABA: Edited MR spectroscopy at 24 research sites. NeuroImage. 2017;159:32–45. doi:10.1016/j.neuroimage.2017.07.021

16. Mikkelsen M, Rimbault DL, Barker PB, et al. Big GABA II: Water-referenced edited MR spectroscopy at 25 research sites. NeuroImage. 2019;191:537–548. doi:10.1016/j.neuroimage.2019.02.059

17. Craven AR, Bhattacharyya PK, Clarke WT, et al. Comparison of seven modelling algorithms for γ-aminobutyric acid–edited proton magnetic resonance spectroscopy. NMR in Biomedicine. Published online February 23, 2022. doi:10.1002/nbm.4702

18. Lin A, Andronesi O, Bogner W, et al. Minimum Reporting Standards for in vivo Magnetic Resonance Spectroscopy (MRSinMRS): Experts’ consensus recommendations. NMR in Biomedicine. Published online February 9, 2021. doi:10.1002/nbm.4484

19. Klose U. In vivo proton spectroscopy in presence of eddy currents. Magn Reson Med. 1990;14(1):26–30. doi:10.1002/mrm.1910140104

20. Mikkelsen M, Tapper S, Near J, Mostofsky SH, Puts NAJ, Edden RAE. Correcting frequency and phase offsets in MRS data using robust spectral registration. NMR in Biomedicine. 2020;33(10). doi:10.1002/nbm.4368

21. Craven AR, Dwyer G, Ersland L, et al. GABA, glutamatergic dynamics and BOLD contrast assessed concurrently using functional MRS during a cognitive task. NMR in Biomedicine. Published online October 28, 2023:e5065. doi:10.1002/nbm.5065

22. Ernst T, Chang L. Elimination of artifacts in short echo time ^1^ H MR spectroscopy of the frontal lobe. Magnetic Resonance in Med. 1996;36(3):462–468. doi:10.1002/mrm.1910360320

23. The MathWorks Inc. MATLAB version: 9.10.0 (R2021a). Published online 2021. https://www.mathworks.com

24. Craven AR, Bell TK, Ersland L, Harris AD, Hugdahl K, Oeltzschner G. Linewidth-Related Bias in Modelled Concentration Estimates from GABA-Edited ^1^ *H-MRS*. Neuroscience; 2024. doi:10.1101/2024.02.27.582249

25. Maudsley AA. Spectral Lineshape Determination by Self-Deconvolution. Journal of Magnetic Resonance, Series B. 1995;106(1):47–57. doi:10.1006/jmrb.1995.1007

26. Sima DM, Garcia MIO, Poullet J, et al. Lineshape estimation for magnetic resonance spectroscopy (MRS) signals: self-deconvolution revisited. Meas Sci Technol. 2009;20(10):104031. doi:10.1088/0957-0233/20/10/104031

27. De Graaf AA, Van Dijk JE, BoéE WMMJ. Quality: quantification improvement by converting lineshapes to the lorentzian type. Magnetic Resonance in Med. 1990;13(3):343–357. doi:10.1002/mrm.1910130302

28. Metz KR, Lam MM, Webb AG. Reference deconvolution: A simple and effective method for resolution enhancement in nuclear magnetic resonance spectroscopy. Concepts in Magnetic Resonance. 2000;12(1):21–42. 10.1002/(SICI)1099-0534(2000)12:1<21::AID-CMR4>3.0.CO;2-R

29. Bartha R, Drost DJ, Menon RS, Williamson PC. Spectroscopic lineshape correction by QUECC: Combined QUALITY deconvolution and eddy current correction. Magn Reson Med. 2000;44(4):641–645. doi:10.1002/1522-2594(200010)44:4<641::AID-MRM19>3.0.CO;2-G

30. Bottomley PA, Griffiths JR, eds. Handbook of Magnetic Resonance Spectroscopy in Vivo: MRS Theory, Practice and Applications. Wiley; 2016.

31. Abramowitz M, Stegun IA. Handbook of Mathematical Functions: With Formulas, Graphs and Mathematical Tables [Conference under the Auspices of the National Science Foundation and the Massachussetts Institute of Technology]. Unabridged, unaltered and corr. republ. of the 1964 ed. Dover publ; 1972. https://personal.math.ubc.ca/~cbm/aands/

32. Mullins PG. Considerations for event-related gamma-aminobutyric acid functional magnetic resonance spectroscopy. NMR in Biomedicine. Published online July 25, 2024:e5215. doi:10.1002/nbm.5215

33. Oeltzschner G, Zöllner HJ, Hui SCN, et al. Osprey: Open-source processing, reconstruction & estimation of magnetic resonance spectroscopy data. Journal of Neuroscience Methods. 2020;343:108827. doi:10.1016/j.jneumeth.2020.108827

34. Zhang Y, An L, Shen J. Fast computation of full density matrix of multispin systems for spatially localized *in vivo* magnetic resonance spectroscopy. Med Phys. 2017;44(8):4169–4178. doi:10.1002/mp.12375

35. Simpson R, Devenyi GA, Jezzard P, Hennessy TJ, Near J. Advanced processing and simulation of MRS data using the FID appliance (FID-A)—An open source, MATLAB -based toolkit. Magn Reson Med. 2017;77(1):23–33. doi:10.1002/mrm.26091

36. Kaiser LG, Young K, Meyerhoff DJ, Mueller SG, Matson GB. A detailed analysis of localized J-difference GABA editing: theoretical and experimental study at 4 T. NMR Biomed. 2008;21(1):22–32. doi:10.1002/nbm.1150

37. Zhu XH, Chen W. Observed BOLD effects on cerebral metabolite resonances in human visual cortex during visual stimulation: A functional1H MRS study at 4 T. Magn Reson Med. 2001;46(5):841–847. doi:10.1002/mrm.1267

38. McKinney W. Data Structures for Statistical Computing in Python. In: van der Walt S, Millman J, eds.; 2010:56–61. doi:10.25080/Majora-92bf1922-00a

39. Harris CR, Millman KJ, van der Walt SJ, et al. Array programming with NumPy. Nature. 2020;585(7825):357-362. doi:10.1038/s41586-020-2649-2

40. SciPy 1.0 Contributors, Virtanen P, Gommers R, et al. SciPy 1.0: fundamental algorithms for scientific computing in Python. Nat Methods. 2020;17(3):261–272. doi:10.1038/s41592-019-0686-2

41. Hunter JD. Matplotlib: A 2D Graphics Environment. Comput Sci Eng. 2007;9(3):90–95. doi:10.1109/MCSE.2007.55

42. Waskom M. seaborn: statistical data visualization. JOSS. 2021;6(60):3021. doi:10.21105/joss.03021

43. Shapiro SS, Wilk MB. An Analysis of Variance Test for Normality (Complete Samples). Biometrika. 1965;52(3/4):591. doi:10.2307/2333709

44. Fligner MA, Killeen TJ. Distribution-Free Two-Sample Tests for Scale. Journal of the American Statistical Association. 1976;71(353):210–213. doi:10.1080/01621459.1976.10481517

45. Wilcoxon F. Individual Comparisons by Ranking Methods. Biometrics Bulletin. 1945;1(6):80. doi:10.2307/3001968

46. Holm S. A Simple Sequentially Rejective Multiple Test Procedure. Scandinavian Journal of Statistics. 1979;6(2):65–70.

47. Bonferroni CE. Il calcolo delle assicurazioni su gruppi di teste. In: Studi in Onore Del Professore Salvatore Ortu Carboni. Tipografia del Senato; 1935:13–60.

48. Theil H. A rank-invariant method of linear and polynomial regression analysis. Indagationes mathematicae. 1950;12(85):173.

49. Sen PK. Estimates of the Regression Coefficient Based on Kendall’s Tau. Journal of the American Statistical Association. 1968;63(324):1379–1389. doi:10.1080/01621459.1968.10480934

50. Dang X, Peng H, Wang X, Zhang H. Theil-sen estimators in a multiple linear regression model. Olemiss Edu. 2008;2.

51. Brix MK, Ersland L, Hugdahl K, et al. Within- and between-session reproducibility of GABA measurements with MR spectroscopy: Reproducibility of MRS GABA Measurements. J Magn Reson Imaging. 2017;46(2):421–430. doi:10.1002/jmri.25588

52. Tapper S, Tisell A, Helms G, Lundberg P. Retrospective artifact elimination in MEGA-PRESS using a correlation approach. Magn Reson Med. 2019;81(4):2223–2237. doi:10.1002/mrm.27590

53. Tal A. The future is 2D : SPECTRAL-TEMPORAL fitting of dynamic MRS data provides exponential gains in precision over conventional approaches. Magnetic Resonance in Med. 2023;89(2):499–507. doi:10.1002/mrm.29456

54. Clarke WT, Ligneul C, Cottaar M, Ip IB, Jbabdi S. Universal dynamic fitting of magnetic resonance spectroscopy. Magnetic Resonance in Med. 2024;91(6):2229–2246. doi:10.1002/mrm.30001

55. Chong DGQ, Kreis R, Bolliger CS, Boesch C, Slotboom J. Two-dimensional linear-combination model fitting of magnetic resonance spectra to define the macromolecule baseline using FiTAID, a Fitting Tool for Arrays of Interrelated Datasets. Magn Reson Mater Phy. 2011;24(3):147-164. doi:10.1007/s10334-011-0246-y

56. Zöllner HJ, Davies-Jenkins C, Simicic D, Tal A, Sulam J, Oeltzschner G. Simultaneous multi-transient linear-combination modeling of MRS data improves uncertainty estimation. Magnetic Resonance in Med. 2024;92(3):916–925. doi:10.1002/mrm.30110

57. Simicic D, Zöllner HJ, Davies-Jenkins CW, Hupfeld KE, Edden RAE, Oeltzschner G. Model-Based Frequency-and-Phase Correction of 1H MRS Data with 2D Linear-Combination Modeling. Neuroscience; 2024. doi:10.1101/2024.03.26.586804

